# *Plantago lanceolata* and *Lolium perenne* metabolite profiles, their impact on soil microbial community structures and soil biological nitrification inhibition

**DOI:** 10.64898/2026.08.17.745343

**Authors:** M. Peterson, N. Joyce, J. van Klink, P. Panda, T. Fraser, C. Anderson

## Abstract

**Background and aims:** Excess nitrate (NO_3_^-^), from fertilizer overuse and intensive agriculture, can pollute water and contribute to greenhouse gas production (nitrous oxide - N_2_O). Plant metabolites from pastural herbs such as *Plantago lanceolata* (plantain) can inhibit microbial nitrification of ammonium to NO_3_^-^ (biological nitrification inhibition - BNI) and change soil nitrogen cycle dynamics (lower potential nitrification rate - PNR). The main aim was to investigate differential plant metabolite expression associated with BNI and lowered PNR in different soil types.

**Methods:** Six plantain cultivars were tested for BNI potential and screened for metabolites that correlated with inhibition of the ammonia oxidising bacterium (AOB) *Nitrosospira multiformis*. PNR and microbiome change was then investigated in four different New Zealand soils under the plantain cultivar ‘Agritonic’ and ryegrass cultivar ‘One50’.

**Results:** PNR under plantain was 11 to 41% lower than fallow soil while PNR under ryegrass was 0 to 39% lower. In addition to verbascoside and aucubin, plantain metabolites associated with lower PNR included plantamajoside, riboflavin 3- and 5-sulfate, plantagoguanidinic acid. Chlorogenic acid was associated with lowered PNR under ryegrass. PNR reductions, microbiome structure and the ratio of ammonia oxidising archaea (AOA) relative to AOB was modulated by soil type.

**Conclusion:** Plantain and ryegrass lowered the PNR in four different soils and was correlated with metabolites beyond just aucubin and verbascoside. Based on candidate BNI-associated metabolites identified, it was hypothesised that lowered PNR is likely indirect through mechanisms such as chelation and appears to be dependent on both plant physiology and soil physicochemistry.

## Introduction

In intensively managed agroecosystems, there is increasing concern regarding the adverse effects of leached nitrate (NO_3_^-^) on water quality and gaseous nitrogen (N) emissions on the atmosphere (Cameron et al. 2013). Soil N dynamics are modulated through complex biochemical interactions between plants, animals, and microbes. Historically, soil N-cycling was thought to be solely under microbial control, but contemporary research has shown that plant-derived compounds can affect microbial N-metabolism both directly or indirectly when exuded into the rhizosphere (Ishikawa et al. 2003; Huberty et al. 2020; Huberty et al. 2022; Kuppe and Postma 2024). These plant compounds are referred to as biological nitrification inhibitors (BNI) because they naturally inhibit microbially mediated nitrification, delaying the conversion of ammonium (NH_4_^+^) to nitrite (NO_2_^-^) and subsequently, NO_3_^-^ production and/or N oxide emissions. It should also be noted that the hydrolysis of urea to NH_4_^+^ by urease can also be affected by plant metabolites thereby impacting down-stream nitrification processes that would also be observed as BNI (Rana et al. 2021).

The use of synthetic nitrification inhibitors (SNI) such as nitrapyrin, 3,4-dimethylpyrazole phosphate and dicyandiamide have been shown to reduce NO_3_^-^ leaching, N_2_O emissions and NO emissions by 48%, 44% and 24% respectively (Qiao et al. 2015). However, their use is limited by cost, application difficulties, a short effective half-life in soil, potential plant damage and concerns over carry-over into food products (Macadam et al. 2003; Fillery 2007; Astley 2013; Zou et al. 2014).

The *in-situ* exudation of BNI by plant roots is a potential low-cost, continuous and targeted alternative to SNI. Several plant species have been identified that exude compounds from their roots to inhibit soil nitrification. Tropical pasture grasses (e.g. *Brachiaria humidicola*), field crops (e.g. *Sorghum bicolor*) and weeds (e.g. *Raphanus raphanistrum*) have been shown to reduce soil nitrification through the root exudation of BNI (Subbarao et al. 2009; Subbarao et al. 2013; O’Sullivan et al. 2017b). The BNI compounds isolated from root exudates and plant tissues include a diverse range of metabolites, including phenolic acids (Rice and Pancholy 1974), fatty acids (Subbarao et al. 2008), phenylpropanoids (Zakir et al. 2008), flavonoids (Subbarao et al. 2013), quinones (Tesfamariam et al. 2014), diterpenoids (Subbarao et al. 2009), monoterpenes (Ward et al. 1997) and isothiocyanates (Brown and Morra 2009).

The genus Plantago (Plantaginaceae) is known to contain a suite of plant secondary metabolites (PSM) that belong to several of the chemical classes noted above (Goncalves and Romano 2016). In Plantago species, the iridoid (monoterpenoid) glucosides aucubin and its derivative catalpol, along with the phenylethanoid glycoside verbascoside (acteoside) have been identified as biologically active (Dietz et al. 2013), including having antimicrobial (Rumball et al. 1997) and antifungal properties (Oyourou et al. 2013).

Aucubin and verbascoside occur in ribwort plantain (*Plantago lanceolata*) at relatively high concentrations (up to 3 and 9% of dry matter respectively), depending on cultivar, plant age, nutrient supply, temperature and light quality (Bowers and Stamp 1993; Tamura 2002; Box et al. 2019) and there is evidence to suggest that these compounds may act as BNIs in soil. Rauber et al. (2008) observed a decline in N mineralisation in field plots when potatoes were under-sown with *P. lanceolata*, while soil mesocosms dominated by *P. lanceolata* had a significantly lower NO_3_^-^concentration, mineralisation and nitrification rates compared to mesocosms dominated by sweet vernal grass (*Anthoxanthum odoratum)* or common bird’s-foot trefoil (*Lotus corniculatus)* (Massaccesi et al. 2015). Additionally, a soil incubation study demonstrated the suppression of N mineralisation and nitrification following incorporation of *P. lanceolata* leaf material into soil (Dietz et al. 2013) while the incorporation of *P. lanceolata* cv. *‘*Tonic’ (percusor to cv. ‘Agritonic’ used in this research) into New Zealand pasture swards has been observed to reduce N-leaching under urine patches (Carlton et al. 2019). Substantial reductions in N leaching >45% have been observed when *P. lanceolata* is present in swards (Woods et al. 2018; Egan et al. 2025; Healy et al. 2026).

Addition of *P. lanceolata* has also been shown to reduce nitrous oxide (N_2_O) emissions from urine patches where the urine was obtained from dairy cows fed a ryegrass/clover diet (Simon et al. 2019). Simon et al (2019) postulate that this emission reduction is a *P. lanceolata* root exudate related BNI effect. In a related study using the same soil type, Simon et al. (2021) also observed a decline in the gene copies of ammonia mono-oxygenase (*amo*A), the translated enzyme of which oxidises NH_4_^+^ to NO_3_^-^, suggesting a linkage between plantain rhizosphere chemistry and BNI. Other studies question the ability of plantain to reduce N_2_O emissions with sward age, soil type and below ground nutrient partitioning cited as reasons for the range of emission results observed including elevated emissions under plantain (Rodriguez et al. 2020; Vi et al. 2023; Pinxterhuis et al. 2024; Ding et al. 2025; Morton et al. 2025; Hammond et al. 2026). However, it should be noted that soil physical conditions such as moisture and temperature have a large influence on N partitioning, soil microbial community N metabolism and likely N loss pathways (Bracken et al. 2022).

Aucubin, catalpol and verbascoside are clearly important with respect to BNI but have not been confirmed as directly responsible for the BNI observed in soil under ribwort plantain. Although multiple studies measure these metabolites in *P. lanceolata* biomass, exudation of these compounds into the rhizosphere is presumed, with BNI activity assumed based on root extractions or addition of leaf material containing these BNI-associated compounds. There is little knowledge about attenuation, mobility and metabolism of exudate compounds once they are released into soil, nor whether there is soil type and mixed sward effects and how that might impact efficacy of BNI expressing plants. There is evidence that microbial community metabolism can be altered by exudate metabolites, especially phenolic compounds (Zhou et al. 2025), but more knowledge about *in situ* soil microbiome behaviour is still required, especially across different soil types. The aggregate inhibition of microbial species with differing sensitivity to BNI compounds (Kaur-Bhambra et al. 2022; Kolovou et al. 2023) will lead to wide variation in the potential nitrification rates (PNR) expressed, especially within rhizosphere soil.

The experiments presented here sought to achieve three aims. The first was to screen plantain cultivars to ascertain the presence of aucubin, catalpol and verbascoside in the above- and below-ground plantain biomass and assess any associated BNI activity. The second aim was to identify other BNI associated metabolites, especially those exuded into soil and postulate on expression pathways. The third aim was to investigate sources of BNI/PNR variability, especially related to soil type and rhizosphere microbiome change.

To meet these aims a preliminary screen of five plantain cultivars and one commercially promoted plantain cultivar was conducted focussing on BNI capacity and shared metabolites. Two cultivars that covered a wide range of BNI capacity were then carried forward, along with the commercial cultivar for a more in-depth comparison of their metabolomes. Perennial ryegrass (*Lolium perrene*) was included as a comparative plant species because it has potential BNI (Sessoms, 2022) and is common in mixed pasture swards. The commercially promoted plantain cultivar and ryegrass were then grown in four different soil types to investigate any differences in root exudate impact on PNR and microbial community response. Our overall hypothesis for this work was that differential expression of BNI associated metabolites delineates BNI capacity and PNR variability across different soil types.

## Materials and Methods

### Root exudate collection from plantain grown in hydroponics and assessment of BNI

Plantain (*Plantago lanceolata* cv. ‘Agritonic’ and several breeding lines designated B-F) cultivars were supplied as seed by Agricom, New Zealand. Approximately 40 seeds of each cultivar were arranged on germination paper (Hoffman Manufacturing Inc., Corvallis, Oregon, USA) in Petrie plates on the laboratory bench under natural light and at ambient temperature (20°C). The germination paper was soaked in deionised water to imbibe the seeds, and water was added as required over the course of germination. Once of sufficient size and vigour, seedlings were transplanted into 4-cm diameter hydroponic baskets, filled with baked clay pellets (CANNA, Subiaco, WA, Australia) to support the plant roots in solution in a hydroponic system.

Twenty-litre plastic boxes were modified into discrete hydroponic growth containers with the capacity to grow nine plants of each cultivar per container. Each container of nine plants represented one replicate, and there were four replicates for each cultivar. These containers were housed in an insulated grow room at a constant temperature of 19 ± 1 °C. The containers were arranged in a spatially adjusted block design under Heliospectra LX601C LED grow lights (Heliospectra AB, Gothenburg, Sweden) on a 16-h light:8-h dark cycle. The photosynthetically active radiation (PAR) averaged 1000 µmol m^-2^ s^-1^ across the hydroponic system when the lights were at full intensity. The hydroponic nutrient solution contained Ca(NO_3_)_2_.4H_2_O (90 mg L^-1^), (NH_4_)_2_SO_4_ (50 mg L^-1^), H_3_PO_4_ (50 mg L^-1^), KCl (56 mg L^-1^), CaCl_2_ (55 mg L^-1^), MgSO_4_.7H_2_O (62 mg L^-1^), MnCl_2_.4H_2_O (0.6 mg L^-1^), ZnCl_2_ (0.4 mg L^-1^), H_3_BO_3_ (1.2 mg L^-1^), CuSO_4_.5H_2_O (0.5 mg L^-1^), Na_2_MoO_4_ (0.01 mg L^-1^) and EDTA Fe(III)Na (15 mg L^-1^). Stock solutions were made up in type II laboratory water (Merck Millipore Elix® Essential 10), with the final medium made up with potable tap water (TDS 130 mg L^-1^, Ca^2+^ 22 mg L^-1^, Na^+^ 12.5 mg L^-1^, Cl^-^ 15.2 mg L^-1^, total alkalinity as CaCO_3_ 63 mg L^-1^, pH 8.0). The pH of the nutrient solution was adjusted to pH 5.7 by the addition of NaOH. The N status, electrical conductivity and pH of the growth media were monitored throughout the experiment and the medium in each container refreshed weekly. Aeration of each hydroponic container was maintained using aquarium air stones and pumps to provide constant air supply.

Forty-five days after germination, the plants were removed from the hydroponic system and the root exudates collected from each plant using a method adapted from that described by Subbarao et al. (2006). After removal from the hydroponic system, the roots were immersed in deionised water (80 mL) and left in the dark for 24 h. After 24 h, the plants were removed from the solution which was then considered the root exudate sample. The exudates collected from the nine plants in each replicate of each cultivar were then combined to represent the exudate profile in that population. Aliquots of each of these combined samples were washed through a Phenomenex StrataX C18 SPE column (2 g/20 mL, Torrance, CA, USA) with two column volumes of type I water (Thermo Scientific™ Barnstead™ Easypure™ II). The adsorbed organic fraction was eluted with 90:10 v/v ethanol:water (20 mL), after which the ethanol and water were driven off by Speedvac-assisted evaporation (Labconco, Kansas City, KS, USA) and the lyophilised exudates weighed and stored at -20°C, pending an assessment for BNI. This concentration step was repeated for each sample to provide an additional lyophilised sample for metabolomic profiling.

Lyophilised exudates were resuspended at a concentration of 1 mg mL^-1^ in a mineral broth (Watson and Mandel 1971) and tested for BNI activity using the bioassay described by (O’Sullivan et al. 2017a). Here, nitrification inhibition by the root exudates was assessed in a plate-based bioassay where Griess reagent (sulfanilamide and N-1-naphthylethylenediamine dihydrochloride under acidic -phosphoric acid - conditions) was used to monitor nitrite (NO_2_^-^) production in pure cultures of the nitrifying bacterium *Nitrosospira multiformis* (ATCC® 25196™) as it metabolises ammonium (NH_4_^+^). The rate of nitrification in the presence of each root exudate was calculated from a linear regression of NO_2_^-^ formation measured every 15 min for 1 h and the BNI activity of the root exudates was calculated by determining the percentage decrease in nitrification rate in the assay containing root exudate relative to uninhibited controls. Statistical tests were performed using data analysis tools in MS Excel, and the k-means cluster analysis with Chi-squared test were performed in R (R Core Team 2026).

### Metabolomic profiling of plantain metabolites

After root exudate collection, the leaf and root of each plant were separated from one another at the stem:root interface and those from the same container combined to represent the population. Both biomass components were frozen at −80°C then freeze-dried until completely dry, taking care, particularly for plantain, that there was no remaining moisture around the corm. The combined samples were weighed then ground through a screen (1mm) using a Pulverisette 15 cutting mill (Fritsch GmbH, Idar-Oberstein, Germany). The ground samples were extracted using methods described by Suomi et al. (2000) and (Stanisavljević et al. 2008). Briefly, plant tissue (c. 5 mg) was weighed into a micro-tube (2 mL) containing 70:30 ethanol:water (1000 μL) and left overnight at a temperature of −20°C. Samples were removed from the freezer and each tube vortex-mixed and homogenised using ultrasound at 37 Hz for 30 min. Following centrifugation (13000 × g) an aliquot (400 µL) of supernatant was filtered using a Thompson™ Single Step® LC filter vial with 0.22 μm PVDF filter. A ‘quality control multi-mix’ of each sample group was prepared by combining an equal aliquot (20 µL) from each sample into a vial. A blank was prepared using the extraction solvents and included in the metabolomic analyses described below.

Plant material and exudates were submitted for ultra-high performance liquid chromatography mass spectrometry (UHPLC-MS) using targeted analysis to quantify concentrations of the known bioactive compounds (aucubin, catalpol and verbascoside) and untargeted metabolomic profiling to discriminate other compounds of interest.

The system consisted of a Thermo Scientific™ (San Jose, CA, USA) Q Exactive™ Plus Orbitrap coupled with a Vanquish™ UHPLC system (Binary Pump H, Split Sampler HT, Dual Oven), calibrated prior to each sample analysis batch with Thermo™ premixed solutions (Pierce™ LTQ ESI Positive and negative ion calibration solutions, catalogue numbers: 88322 and 88324, respectively).

An aliquot (2 μL) of each sample preparation was separated with a mobile phase consisting of 0.1% formic acid in type 1 water (A) and 0.1% formic acid in acetonitrile (B) by reverse phase chromatography (Hypersil GOLD™ aQ C18 1.9 µm, 100 mm × 2.1 mm, P/N: 25302-102130, Thermo Scientific) maintained at 40°C with a flow rate of 300 µL min^-1^. A gradient was applied: 0–1 min/0% B, linear increase to 2 min/20% B, linear increase to 9 min/45% B, linear increase to 10 min/98% B, isocratic to 13 min/98% B, linear decline to 14 min/0% B, isocratic to end 17 min/0% B.

The eluent was scanned from 0.7 to 12 min respectively by API-MS (Orbitrap) with heated electrospray ionisation (HESI) at 350°C in the negative and positive mode with capillary temperature of 320°C. Data were acquired for precursor masses *m/z* 110–1500 amu at 70K resolution (AGC target 3e6, maximum IT 100 ms, profile mode) with data-dependent ms/ms for product ions generated by normalised collision energy ((C) NCE:15, 35, 85 and at 17.5K resolution (TopN 10, AGC target (C) 1e5 (H) 2e5, Maximum IT 50 ms, Isolation 1.4 *m/z*).

Datasets were then processed with the aid of Compound Discoverer 3.3.3.200 (Thermo Fisher Scientific) for quality control and discriminant analysis. The context to align batch files and compound features was achieved by importing an experimental design study file array providing identifiers for processing of analytical input files as ‘blanks’, ‘sample’, ‘standard’, and ‘quality control’ with the grouping context of experimental ‘categorical’, ‘biological’ and ‘numerical’ factors (i.e. leaf, root, cultivar, soil type, assay value).

Chromatographic and spectrometry data were filtered by a node based process: 1) ‘peak rating’ of 5.5 (scale 0-10) in at least 5 samples, 2) ‘similar features search’ within a tolerance of 5 ppm mass tolerance, 3) ‘Centroids Filtering’ with S/N threshold = 1.5, 4) ‘Real peak detection’ (for more accurate areas) was performed, 5) ‘QC correction’ performed by using SERRF method (Fan et al. 2019), 6) ‘Random Forest’ was run with 200 trees, 7) Differential analysis used peak area with data transformation log-10 areas for p-value estimation with the p-value of per group ratio calculated by a one-way ANOVA model with Tukey as post-hoc test with correction p-value adjusted using Benjamini-Hochberg correction for the false-discovery rate; p-value set at < 0.05 and log2fold change >1.

### PNR as a proxy for BNI under plantain and ryegrass grown in different soil types

Soils for determining the effect of root exudation on nitrification were collected from the Plantain Potency and Practice research programme National Plot Trial sites across New Zealand. A gley (Aquic) and allophanic (Andisol) were collected from the Waikato site (Latitude - 37.9157, Longitude 175.1926) while pallic soils were collected from Manawatū (-40.1332, 175.6545) and Canterbury (-43.6124, 172.4688) sites. At each sampling site, soil was collected from buffer zones that were under permanent ryegrass, to a depth of 10 cm (total ∼20 kg soil collected). The soils were sieved to 6 mm while field moist, then air-dried. Subsamples were sieved to 4 mm and 2 mm and were analysed for chemical and physical characteristics (Supplementary Tables 1-4), with methods detailed below.

Soils were packed to 80% of the dry bulk density into two-compartment pots (rhizopots) consisting of a PVC planter pot (80 mm height, 75 mm Ø) and a buffer pot (40 mm height, 75 mm Ø). Twenty-micron mesh (Sefar Nitex 03-20/14) separated the two compartments while polyester voile was used to retain the soil in the buffer pot. The buffer pots were arranged in a spatially adjusted block design on a sand table (Supplementary Figure 1). The sand table was flooded with water and the buffer pots allowed to equilibrate to field capacity before the planter pots were stacked on top and the two parts taped together to form the complete rhizopot. The complete rhizopots were left for 7 days to equilibrate to field capacity before sowing.

Seeds of ‘Agritonic’ plantain (*Plantain lanceolata*) and ‘One50’ ryegrass (*Lolium perenne*) (Agricom, New Zealand) were sown at a rate of 10 seeds per pot, thinned to 3 after germination. There were seven pots per soil type x treatment, with the third treatment being unplanted (fallow) pots. The sand table on which these pots were arranged was in a glasshouse. The photoperiod at the start of the experiment was 14 h day:10 h night; this was maintained for the duration of the experiment using supplementary lighting. The glasshouse was automatically vented so that average daily temperatures did not exceed 25°C and heaters were deployed when the temperature dropped below 12°C. The plants were left to grow for 90 d to allow the roots to explore the full volume of the pots. Soil moisture was maintained at field capacity using a flood irrigation system. Fertiliser was applied to the pots 3 weeks post-germination. Phosphorus, potassium, and sulfur (as K_2_SO_4_ and KH_2_PO_4_) were applied elementally at rates of 40, 100 and 20 mg kg^-1^ soil, respectively, to ensure that none were limiting (Curtin and McCallum 2004). Eight weeks post-germination, 50 mg N kg^-1^ in the form of liquid urea was added to all pots.

At harvest, the plants were removed from the pots, and soil was shaken from the roots into plastic bags. The moisture content of the soils was determined and the potential BNI effect in soils was assessed using the shaken-slurry potential nitrification rate (PNR) assay (Drury et al. 2008) where a lowering of PNR was presumed to be predominantly related to a BNI effect from exuded root metabolites. Briefly, 15 g soil was placed in 100 mL buffered medium containing 1 mM PO_4_^3-^ and 1.5 mM NH_4_^+^ (pH ∼7.2) and incubated at 20°C on a shaking platform rotating at 180 rpm. Samples of the slurry were collected at 0, 2, 6, 20 and 24 h after the incubation was started, and the liquid fraction was analysed for NO_3_^-^ using a Lachat QuikChem 8500 Series 2 Flow Injection Analysis System (Lachat Instruments, Loveland, Colorado, USA). The PNR was determined by linear regression of NO_3_^-^ concentration over the incubation time as described by Weaver et al. (1994).

Data were visualised in GenStat (64-bit Release 22; VSN International Ltd, Hemel Hempstead, UK) and a mixed-effects model (with fixed effects for soil and treatment and random effects for row and column of the layout) used to provide statistical analysis of the data. SigmaPlot v15.0 (Systat Software Inc., DE, USA) graphical software was used.

### Plant nitrogen measurement and metabolite profiling in soil

The above-surface biomass (ASBM) and below-surface biomass (BSBM) of each plant were separated from each other, with the BSBM being washed before both biomass components were frozen at -80°C and freeze dried. The samples were weighed then ground through a 1 mm screen using a Kannastör® GR8TR V2 fine grinder (Volo Trade Inc., Miami, FL). After grinding, the samples were dried for 16 h at 60°C and total N determined on a 0.1 g sub-sample by automated dry combustion (at 1250°C) using a TruMac CN analyser (Leco Corporation, MI, USA).

Root-associated soil was collected from the plants grown in the pots described above. A minimum of 2.0 g of each soil was sampled aseptically into two separate 2 mL microcentrifuge tubes and frozen at −80°C pending extraction and analysis for metabolomics and DNA extraction. For untargeted metabolomic analysis the soils were extracted using the method of Swenson and Northen (2019). LC-MS grade water (4 mL) was added to soil samples (1.0 g) and sonicated (50% amplitude for 2 × 30 s) and placed on a refrigerated orbital shaker at 200 rpm for 1 h. After centrifugation (3220 × g for 15 min at 4°C) and filtration (0.45 μm PVDF), the extracts were frozen at −80°C before lyophilisation. The lyophilised samples were resuspended in LC-MS grade methanol, vortexed, centrifuged (5000 × g for 5 min) and an aliquot of resuspended extract transferred to a Single Step® 0.22-μm PVDF filter vial (Thompson™ Part No. 65531-200) for LC-MS analysis and feature filtration and curation and analysis as described above.

### Soil chemistry analysis

Soil pH was analysed in a 1:2.5 air-dried, <4 mm sieved soil to water slurry (or field-moist equivalent), vigorously stirred then left to stand for 1 h before measurement with a soil specific combination pH electrode (Devey et al. 2010). Soil conductivity was measured in a 1:5 air-dried soil to water suspension, shaken for 1 h then left to stand for 1 h before measurement with a conductivity probe (Rayment and Lyons 2011).

Olsen P was measured following a temperature-controlled extraction at 25°C of 2.0 g of air-dried soil, (sieved to <2 mm) in 40 mL of 0.5 N NaHCO_3_ solution buffered to pH 8.50 +/- 0.05 with end-over-end tumbling for 30 min (Olsen et al. 1954). The extracts were centrifuged and filtered immediately after tumbling and analysed for orthophosphate by flow injection analysis.

Total carbon and nitrogen were determined on soil sieved to <2 mm and oven-dried for 16 h at 60°C. The samples were dry combusted (Dumas) using TruMac CN analyser (Leco Corporation, MI, USA) operating at 1250°C. Carbon was detected as carbon dioxide by non-dispersive infrared absorption and nitrogen by a thermal conductivity detector following scrubbing of carbon dioxide and oxygen.

Mineral N concentrations were determined by 1 h extraction of 5 g field-moist soil (sieved to <4 mm), shaken with 25 mL of 2 M KCl (Keeney and Nelson 1982). The extracts were then centrifuged, filtered, and analysed for inorganic nitrogen (NH_4_-N and NO_x_-N) by flow injection analysis.

Total extractable organic N was determined using a method adapted from Ghani et al. (2003), and further developed by Curtin et al. (2017). Four grams of air-dried soil was shaken for 30 min with 40 mL of cold (room temperature) water, then placed in an 80°C water bath for 16 h before centrifugation and filtration. Inorganic nitrogen (NO_x_-N and NH_4_-N) was measured by flow injection analysis. Total persulfate nitrogen was measured as NO_x_-N, following a digestion of the extract with a persulfate oxidising reagent (1:1) for 16 h at 90°C. Total extractable organic nitrogen was determined by subtracting inorganic N from total persulfate N.

Sulfate-S was measured from a calcium phosphate extract of the soil using an ion-chromatography (Searle 1988). Organic-S was calculated by determining total extractable sulfur on the same extract by ICP-OES and subtracting sulfate-S (Watkinson and Perrott 1990).

Mehlich 3-extractable nutrients were determined by shaking a volume of Mehlich 3 extract, pH 2.6 (containing ammonium fluoride, ammonium nitrate, EDTA and acetic and nitric acids) with soil followed by clarification then analysis by ICP-OES (Mehlich 1984).

Total extractable nutrients were determined by refluxing soils with aqua regia (hydrochloric and nitric acids). After extraction, the solubilised analytes were diluted to specified volumes with ASTM Type I water either centrifuged or allowed to settle overnight before analysis by ICP-OES.

Microbial biomass N was determined using the chloroform fumigation extraction technique (Vance et al. 1987). Ten grams of soil (field-moist, sieved to <4 mm) was fumigated in the dark for 48 h then extracted with 40 mL of 0.5 M K_2_SO_4_, with a second 10 g sample extracted without fumigation. Total organic nitrogen in the extracts was measured as NO_x_-N by flow injection analysis, following a digestion of the extract with a persulphate oxidising reagent (1:1) for 16 h at 90°C (Cabrera and Beare 1993). Microbial biomass N was calculated from the difference between the fumigated and non-fumigated samples.

Statistical analysis and tests were performed using data analysis tools in MS Excel with any multivariate analysis conducted in PRIMER-E with PERMANOVA add-on (PRIMER 7, PRIMER-E Ltd).

### Assessing soil microbial community change

DNA from frozen soil subsamples (2 g) was extracted with the DNeasy PowerLyzer Powersoil® DNA kit (Qiagen) as per the manufacturer’s instructions. DNA quality and quantity was assessed by UV absorbance (NanoDrop ND-1000, ThermoFisher Scientific) with most samples having a 260/280 ratio of 1.8–2.0. DNA concentrations varied between 8.5 ng mL^-1^ and 103.1 ng mL^-1^, with an average of 50.2 ng mL^-1^.

The V3-V4 variable region of the 16S rRNA gene was amplified using the bacterial-specific 341F (5’-CCTACGGGNGGCWGCAG-3’) and 785R (5’-GACTACHVGGGTATCTAATCC-3’) primer pair (Klindworth et al. 2013). PCR amplifications for the bacterial samples contained 2 μL of 1/10 dilution of template DNA, 1.25 µL of 5 µM forward primer, 1.25 µL of 5 µM reverse primer, 12.5 µL of PCR master mix (MyFi Mix 2X, Bioline), and 8 µL of nuclease-free ddH_2_0 in a final volume of 25 μL. Reactions were performed in duplicate. Cycling parameters were 95°C for 1 min; 25 cycles of 95°C for 30 s, 50°C for 30 s, 72°C for 30 s then 72°C for 5 min.

The archaeal community was characterised from each sample by amplification of the V4-V5 region of the 16S rRNA gene using the archaeon-specific primer pair 516F (5’-TGYCAGCCGCCGCGGTAAHACCVGC-3’) and 915R (5’-GTGCTCCCCCGCCAATTCCT-3’) (Raymann et al. 2017). The PCR mixture for the archaeal samples contained 2 µL of 1/10 dilution of template DNA, 1.25 µL of 5 µM forward primer, 1.25 µL of 5 µM reverse primer, 12.5 µL of PCR master mix (MyFi Mix 2X, Bioline), and 8 µL of nuclease-free ddH20 in a final volume of 25 μL. Cycling parameters were 95°C for 1 min; 35 cycles of 95°C for 20 s, 50°C for 20 s, 72°C for 35 s then 72°C for 5 min.

All primers included the Illumina adapter sequences for the forward overhang (5’ TCGTCGGCAGCGTCAGATGTGTATAAGAGACAG-[locus-specific sequence]-3’) and reverse overhang (5’ GTCTCGTGGGCTCGGAGATGTGTATAAGAGACAG-[locus-specific sequence]-3’).

Duplicate reactions were combined and purified with AMPure XP beads (Agencourt, Beckman Coulter Life Sciences). Purified amplicons were quantified by Quant-iT™ PicoGreen™ dsDNA Assay Kit (Invitrogen, ThermoFisher Scientific). Concentration of amplicons ranged from 3.0 ng μL^-1^ to 60.3 ng μL^-1^ for bacteria and from 3 ng μL^-1^ to 108.1 ng μL^-1^ for Archaea and were normalised to 3 ng μL^-1^ before sending for sequencing. Amplicons were 2 × 300 bp paired end sequenced on an Illumina MiSeq platform (Massey Genome Service, Palmerston North, New Zealand).

Demultiplexed raw reads obtained from Massey were quality-filtered, denoised, chimera-checked and processed into amplicon sequence variants (ASV) using the DADA2 (v1.16.0) R package (Callahan et al. 2016). The 16S rRNA primer sequences were removed using the trimLeft function and truncated at 260 bp (variable, depending on run quality/length) for the forward and reverse reads. The taxonomic assignment of the ASVs was conducted using the assignTaxonomy function in DADA2. The SILVA (v138.1) rRNA database (Quast et al. 2013) was used as the reference database for both bacteria and archaea. Sequences belonging to mitochondria and chloroplasts were removed from the dataset.

Multivariate analysis was conducted in PRIMER-E with PERMANOVA add-on, generally with default parameters throughout (PRIMER 7, PRIMER-E Ltd.).

### Assessment of nitrifier communities using the ammonia monooxygenase (*amoA*) gene

DNA samples were also screened for the gene encoding the alpha-subunit of *amoA* using a duplex real-time quantitative polymerase chain reaction (qPCR) assay combining both the internal control primers and *amoA* primers.

Primers for the 16S rRNA gene were as above without the MiSeq adaptor sequence while the primers specific to the bacterial *amo*A gene used were amoA-1F (5’-GGGGHTTYTACTGGTGGT [H = not G; Y = C or T]) and amoA-2R (5’-CCCCTCKGSAAAGCCTTCTTC [K = G or T; S = G or C]) (Stephen et al. 1999). Archaeal *amo*A primers were as per those outlined in Pereira et al. (2020). The reaction conditions used for the 16S rRNA gene-specific primers were 95°C for 2 min for 1 cycle; and 95°C for 5 s, followed by 55°C for 30 s for 40 cycles. The conditions used for the *amo*A gene-specific primers were identical, except that the annealing temperature was 56°C. For 17 out of 84 samples, an extension step of 72°C per 30 seconds was included after annealing, as Cqs for both sets of primers were too high (30+) in the 2-step reaction. There was no major difference between the 2-step reaction and the 3-step reaction for the rest of the samples, apart from increased qPCR running times.

Each sample contained 2 μL of template DNA, 0.8 μL of forward and reverse primer (5 μM concentration), 5 μL of 2X qPCRBIO SyGreen Blue Mix Hi-ROX (PCR Biosystems), and 1.4 μL of molecular-grade water for a final reaction volume of 10 μL. PCR amplification reactions were performed using a CFX96 real-time PCR detection system (Bio-Rad). Each sample was run in triplicate. Each qPCR included a set of standards and a blank (where the sample was replaced with water). Standards were generated from gDNA extracted from *Nitrosospira multiformis* (ATCC® 25196™) and *Nitrososphaera viennensis* (strain EN76). The qPCRs had an average R2 of 0.96. The average efficiencies of the *amo*A and 16S qPCRs were 101% and 105%, respectively. Melting-curve peaks for the standards and samples amplified using the *amo*A and 16S rRNA gene-specific primers occurred at temperatures ranging from 88.5 to 89.5°C and 86.5 to 87.0°C, respectively, with no other minor peaks being detected indicating the absence of nonspecific binding. Relative differences between samples were compared using delta Cq (*amo*A Cq – 16S rRNA Cq).

Basic statistical analysis and tests were performed using data analysis tools in MS Excel. Multivariate analysis was conducted in PRIMER-E with PERMANOVA add-on, generally with default parameters throughout (PRIMER 7, PRIMER-E Ltd.).

## Results

### BNI and associated plantain metabolite chemistry from hydroponically grown plants

The BNI activity of the root exudates of six plantain cultivars was calculated as the decrease in nitrification rate conferred by a chemically undefined exudate (1mg mL^-1^) relative to that in an uninhibited control. This activity varied considerably between the cultivars and between the replicates of each cultivar (Supplementary Figure 2). The averages for inhibition induced by the exudates from the six cultivars ranged between 22% and 42%. Exudates from Cultivar B generally had lower inhibitory activity while cultivars C, D and ‘Agritonic’ exhibited generally higher inhibitory activity. Although there was an observable trend, the high variability resulted in a lack of statistical support for any differences in inhibition among the plantain cultivars.

A preliminary untargeted LC-MS screening was performed to identify either putative or known metabolites present in the six plantain cultivars. After filtering of background noise, ms/ms spectra threshold and removal of features due to lack of formula prediction, 1544 mass spectral compound features remained from a total of 2507. These 1544 mass features were represented across two modes of instrument source ionisation (+/-) and two chromatography columns (reverse and normal phase) with the metabolite profiles ranging from small organic compounds to polar lipids, including organic acids, amino acids, iridoids, phenolics and terpenes.

The data was then grouped by plantain cultivar and tissue (leaf or root). When constrained by cultivar, correlation of individual metabolite values with BNI assay activity values had poor statistical discrimination, so cultivar independent inhibition ranges were formed instead. The ranges were defined as: ‘low BNI’ (14–23%), ‘medium BNI’ (28–36%) and ‘high BNI’ (41–66%) with these delineations supported by a k-means cluster analysis (Supplementary Figure 3). On this basis, candidate BNI associated metabolites shared between cultivars that had a positive correlation to ‘high BNI’ and negative correlation to ‘low BNI’ were evaluated in both tissue types. Within the ‘high BNI’ range there were at least 39 compounds expressed in the root tissue, with only a few expressing differentially in the leaf. These compounds were largely phenylethanoid glycosides with verbascoside being the largest contributor. Conversely, the iridoid catalpol in the root correlated with the ‘low BNI’ group. Other common compounds included quinic acids such as neochlorogenic acid, along with sugar acids and several unknown compounds requiring further investigation to characterise conclusively.

To lower the dataset complexity and improve discrimination of the putative BNI-associated metabolites discerned in the first hydroponic experiment, a second hydroponic experiment was conducted using ‘Agritonic’, Cultivar E (CE), and Cultivar B (CB) to represent the generalised ‘high’, ‘medium’ and ‘low’ BNI ranges, respectively. These generalised ranges were applied based on the hypothesis that differential expression of metabolites, irrespective of cultivar, was correlated with BNI activity. The first hydroponic experiment indicated that the cultivars selected were collectively capable of at least 14-66% BNI compared to uninhibited controls. Replication per cultivar was increased from four to seven and a ryegrass cultivar (One50) was included as a reference plant species with low expected BNI activity.

After filtering, the compound dataset contained 2633 mass features that distinguished leaf and root metabolites with principal components analysis (PCA) clearly illustrating the difference in leaf versus root metabolites and the separation from ryegrass (Supplementary Figure 4). Further discriminant analysis and filtering of the mass feature data was based on applying the ‘high’, ‘medium’ and ‘low’ BNI ranges as per previous results. Leaf metabolites from volcano plots (*p*<0.05) that distinguished plantain cultivars from ryegrass were dominated by phenylpropanoids and flavins. The major measurable forms detected were caffeoyl phenylethanoid glycosides, where the cinnamic acids are linked via glucose (and associated sugars like rhamnose) to hydroxytyrosol. Although present in high amounts (up to 200 mg g^-1^, phloretic acid equivalent), the bioactive caffeoyl phenylethanoid-phenylpropanoid glycoside target verbascoside, was not indicated as a discriminating biomarker for potential BNI activity for plants grown in the hydroponic system. Instead, the most statistically significant glycoside discriminator was tentatively identified as plantamajoside (Figure 1).

**Figure 1:**
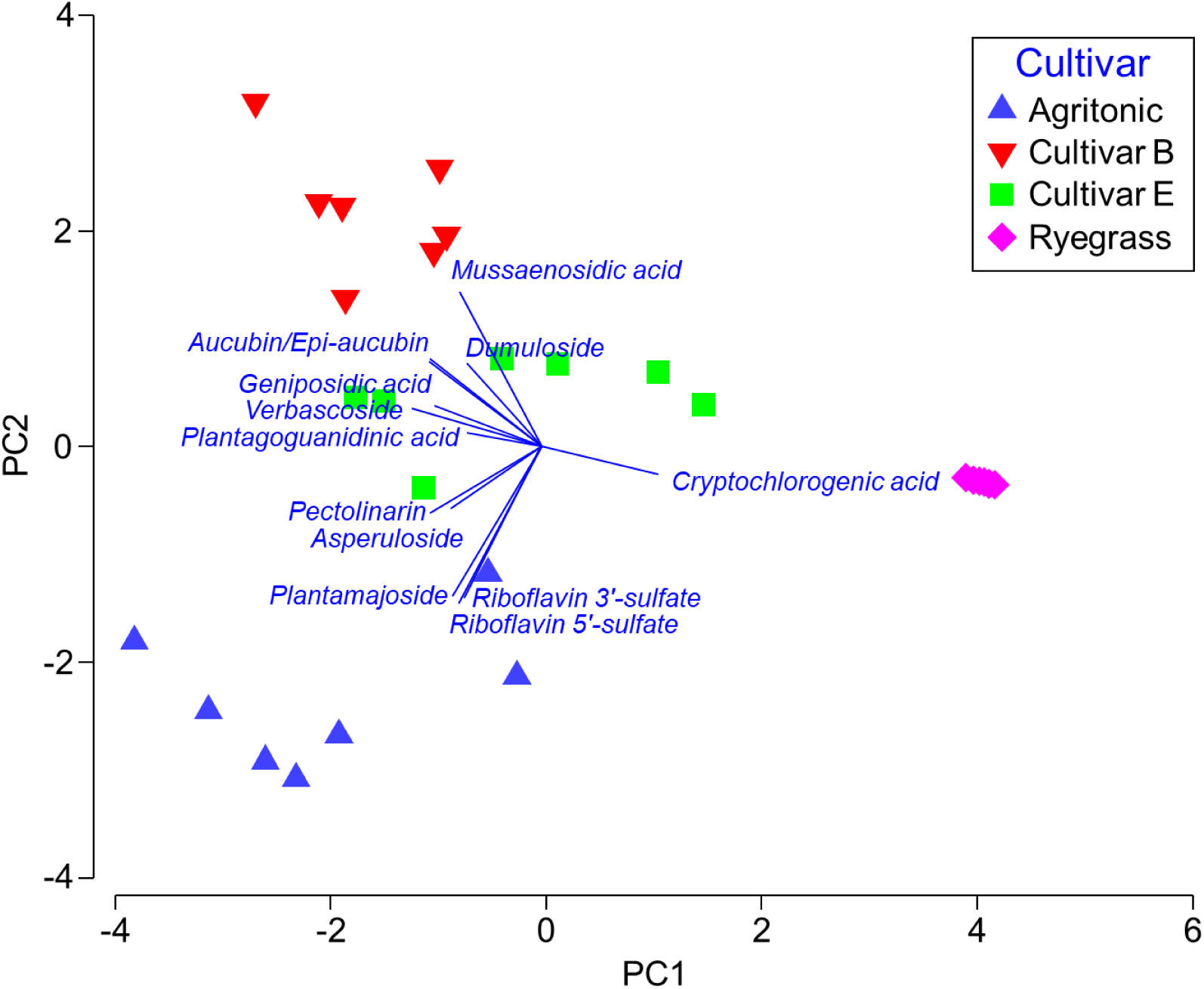
Shared leaf metabolites that are related to BNI activity ranges for plantain cultivars ‘Agritonic’ (High BNI), Cultivar B (low activity) and Cultivar E (medium activity). Ryegrass metabolites captured are those that clearly distinguish this species from plantain. 53.3% of variation is captured by the PC1 axis and 22.6% by PC2. The PC3 axis only captured a further 7.5%. The blue vector fan indicates increasing concentration of each metabolite from the centre out, with length and the direction being an indicator of ‘influence’ that each metabolite has with respect to where data points are positioned in the plot relative to each other.

Discriminating flavin metabolites were tentatively identified as riboflavin-3-sulfate and riboflavin-5-sulfate (Figure 1). The bioactive iridoid target aucubin concentrations did not correlate positively with expected differences in BNI. Instead, asperuloside and pectinolarin were compounds of interest. Aucubin concentrations showed an inverse relationship with respect to expected BNI, along with dumuloside and mussaenosidic acid. Observations for these iridoids in leaves with respect to expected BNI did not equate with those in the roots (Figure 2).

**Figure 2:**
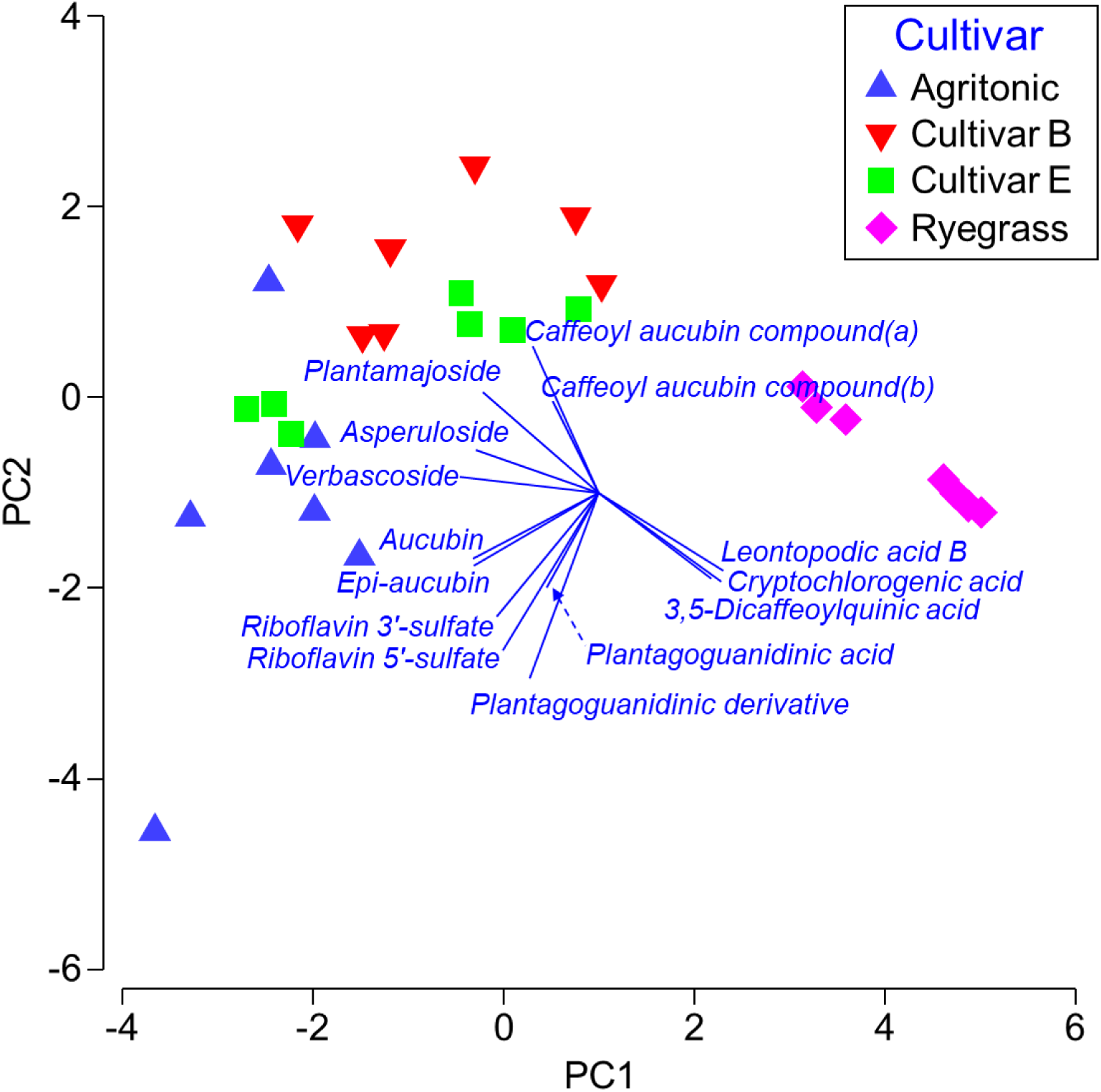
Shared root metabolites that are correlated to BNI activity ranges for plantain cultivars ‘Agritonic’ (High BNI), Cultivar B (low activity) and Cultivar E (medium activity). Ryegrass metabolites captured are those that clearly distinguish this species from plantain. 53.7% of variation is captured by the PC1 axis and 14.3% by PC2. The PC3 axis only captured a further 8.1%. The blue vector fan indicates increasing concentration of each metabolite from the centre out, with length and the direction being an indicator of ‘influence’ that each metabolite has with respect to where data points are positioned in the plot relative to each other.

Several root-expressed compounds distinguished plantain cultivars based on expected BNI, with key metabolites in ‘Agritonic’ root tissue tentatively identified as guanidine derivatives, cinnamic and flavin compounds. These consisted of plantagoguanidinic acid and a related derivative, the two riboflavin sulfates identified in the leaf tissue, and aucubin (Figure 2). Other correlating compounds from the amine-imidazole and flavin classes were also detected in low abundance but require further investigation to characterise conclusively. Ryegrass was characterised by the presence of cryptochlorogenic acid in the leaf tissue, accompanied by isochlorogenic acid and leontopodic acid in the root tissue (Figures 1 and 2).

### Influence of soil type on plantain and ryegrass metabolomes and detectable exudates

The metabolomic profiles of leaf and root tissue from plantain and ryegrass, and their impact on soil nitrification rates, were investigated with respect to growth in different soil types. ‘Agritonic’ was carried forward as the representative plantain cultivar with winter activity, distinctive chemistry and generally ‘high’ BNI activity, and was therefore expected to lower the potential nitrification rate (PNR) in soil. Ryegrass cultivar ‘One50’ was also carried forward.

The effect of active root exudation on PNR in soil varied for both plant species (plantain and ryegrass) and soil type (Figure 3). In this assay, PNR in soil that had been subject to active plant growth was compared with PNR in soil from fallow controls. All soils had low ammonium (NH_4_^+^) levels prior to spiking for the PNR assay with no statistical differences between treatments (Table 1). High background nitrate (NO_3_^-^) concentrations in fallow soils did not appear to impact nitrification with PNR rates easily calculated from NO_3_^-^ accumulation above background. There did not appear to be any relationship between microbial biomass and PNR. Pre-PNR measures of total extractable organic-N and microbial biomass-N were both elevated in soils planted with ryegrass, but these increases were not necessarily statistically supported (Table 1). In the two pallic soils, where differences in microbial biomass-N were not statistically supported, the PNR was lower for both plantain and ryegrass compared to fallow, but where ryegrass exhibited a much higher microbial biomass-N (i.e. the allophanic and gley soils), PNR was not discernible from fallow and plantain was lower (Table 1, Figure 3).

**Figure 3:**
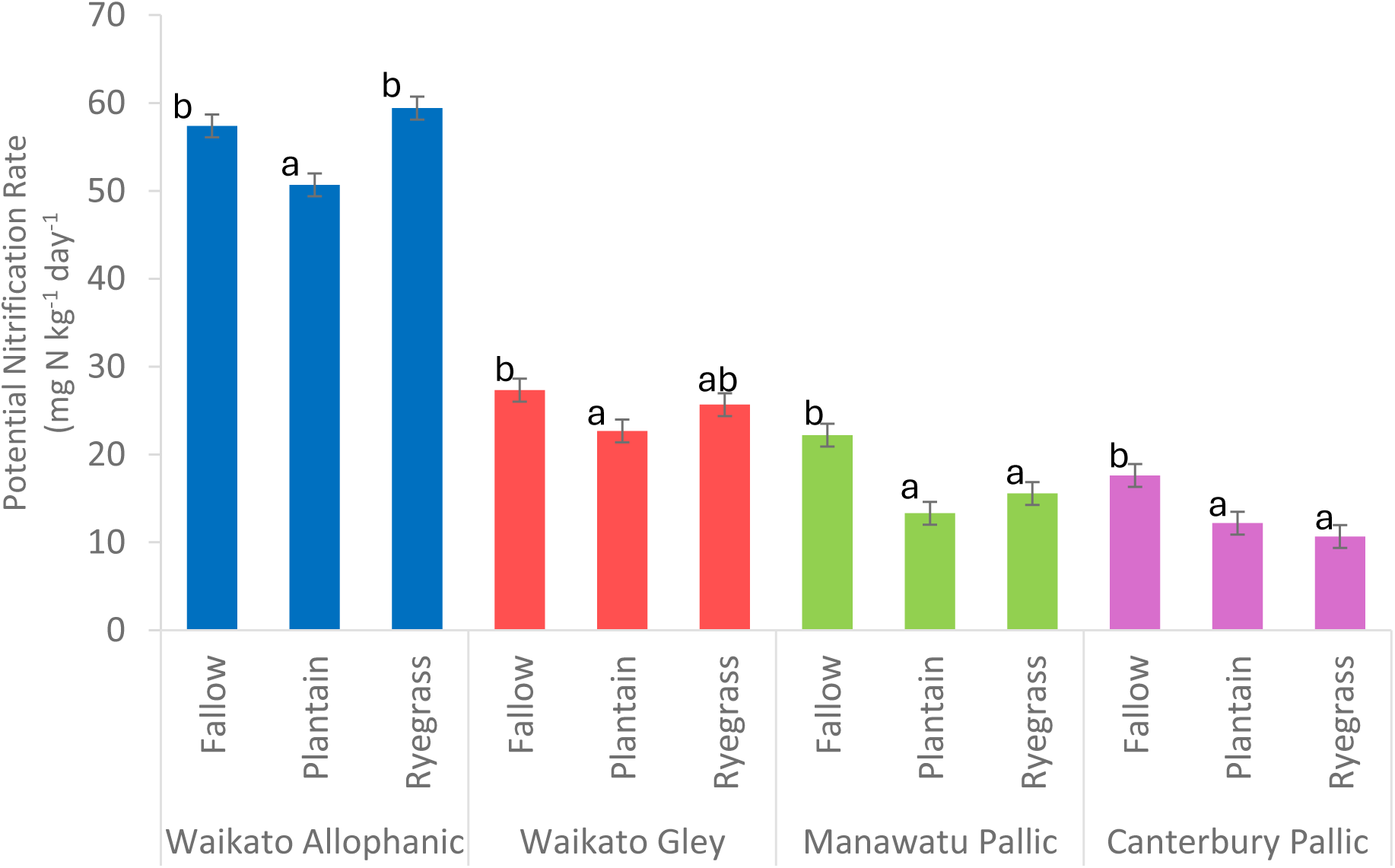
Potential Nitrification Rates (PNRs) in four soils sourced from three New Zealand regions under ‘Agritonic’ plantain and ‘One50’ ryegrass. A letter in common (within a soil) indicates the means are not significantly different, using the least significant difference (*p* < 0.05).

**Table 1:** N partitioning and pH in soil under fallow, plantain or ryegrass treatments prior to the potential nitrification rate assay. Letters after means are comparisons within a soil, using the least significant difference e.g. two means (within a soil) with a letter in common are not significantly different (p < 0.05). Supporting ANOVA can be found in Supplementary Table 5.

|  | pH | Min N<br>(mg N kg<sup>-1</sup>) | NO<sub>x</sub>-N<br>(mg N kg<sup>-1</sup>) | NH<sub>4</sub>-N<br>(mg N kg<sup>-1</sup>) | Total Extractable<br>Organic N<br>(mg N kg<sup>-1</sup>) | Microbial<br>Biomass N<br>(mg N kg<sup>-1</sup>) |
| --- | --- | --- | --- | --- | --- | --- |
| **Waikato Allophanic** |  |  |  |  |  |  |
| Fallow | 5.8 | 168 ± 74<sup>B</sup> | 167 ± 93<sup>B</sup> | 1.4 ± 0.7<sup>A</sup> | 271 ± 5.2<sup>A</sup> | 132 ± 28<sup>AB</sup> |
| Plantain | 6.3 | 4.4 ± 2.0<sup>A</sup> | 1.0 ± 0.6<sup>A</sup> | 3.0 ± 1.4<sup>A</sup> | 248 ± 5.2<sup>B</sup> | 127 ± 7<sup>A</sup> |
| Ryegrass | 6.3 | 2.5 ± 1.1<sup>A</sup> | 0.5 ± 0.3<sup>A</sup> | 1.9 ± 0.9<sup>A</sup> | 286 ± 5.2<sup>C</sup> | 172 ± 4<sup>B</sup> |
| **Waikato Gley** |  |  |  |  |  |  |
| Fallow | 5.6 | 198 ± 87<sup>C</sup> | 196 ± 110<sup>C</sup> | 0.9 ± 0.4<sup>A</sup> | 159 ± 5.2<sup>A</sup> | 34 ± 28<sup>A</sup> |
| Plantain | 6.0 | 5.5 ± 2.4<sup>B</sup> | 3.4 ± 1.9<sup>B</sup> | 1.2 ± 0.6<sup>A</sup> | 159 ± 5.2<sup>A</sup> | 91 ± 7<sup>AB</sup> |
| Ryegrass | 6.5 | 1.5 ± 0.7<sup>A</sup> | 0.4 ± 0.2<sup>A</sup> | 1.1 ± 0.5<sup>A</sup> | 184 ± 5.2<sup>B</sup> | 118 ± 4<sup>B</sup> |
| **Manawatū Pallic** |  |  |  |  |  |  |
| Fallow | 6.2 | 160 ± 70<sup>B</sup> | 155 ± 87<sup>C</sup> | 3.3 ± 1.5<sup>A</sup> | 131 ± 5.2<sup>A</sup> | 44 ± 28<sup>A</sup> |
| Plantain | 6.4 | 2.8 ± 1.2<sup>A</sup> | 0.7 ± 0.4<sup>B</sup> | 1.8 ± 0.8<sup>A</sup> | 136 ± 5.2<sup>A</sup> | 60 ± 7<sup>A</sup> |
| Ryegrass | 6.9 | 1.2 ± 0.5<sup>A</sup> | 0.1 ± 0.1<sup>A</sup> | 1.3 ± 0.6<sup>A</sup> | 154 ± 5.2<sup>B</sup> | 70 ± 4<sup>A</sup> |
| **Canterbury Pallic** |  |  |  |  |  |  |
| Fallow | 5.8 | 94 ± 41<sup>C</sup> | 93 ± 52<sup>C</sup> | 1.8 ± 0.8<sup>A</sup> | 151 ± 5.2<sup>A</sup> | 66 ± 28<sup>A</sup> |
| Plantain | 6.0 | 15 ± 7<sup>B</sup> | 8.0 ± 4.5<sup>B</sup> | 4.4 ± 2.0<sup>A</sup> | 165 ± 5.2<sup>AB</sup> | 54 ± 7<sup>A</sup> |
| Ryegrass | 6.2 | 2.8 ± 1.2<sup>A</sup> | 1.3 ± 0.7<sup>A</sup> | 1.9 ± 0.9<sup>A</sup> | 179 ± 5.2<sup>B</sup> | 68 ± 4<sup>A</sup> |

The PNR in the allophanic soil was 2–3 times greater than in the other soils, with the maximum rate recorded being ∼60 mg N kg^-1^ day^-1^. The PNR in the plantain treatments was lower than in the fallow treatments for all soils; PNR was reduced by 33%, 41%, 15% and 11% for the Canterbury pallic, Manawatū pallic, Waikato gley and Waikato allophanic soils, respectively. Except for the Waikato allophanic soil, PNR in the ryegrass treatments was also substantially reduced by 39% for the Canterbury pallic soil, 27% for the Manawatū pallic soil and to a smaller extent for the Waikato gley soil. The ryegrass associated PNR reductions were not statistically discernible from those under plantain (Figure 3).

As per hydroponically grown plants, verbascoside was a major contributor in the plantain biomass profile, with greater concentrations found in the leaf than the root (up to 21 mg g^-1^ versus up to 15 mg g^-1^, catechin equivalent, respectively) but concentrations were up to 10-fold lower than the hydroponically grown plants. Aucubin was found at concentrations up to 9 and 17 mg g^-1^ in leaf and root tissue respectively - again, lower than hydroponically grown plants, but only up to 5-fold in this case. There was no statistically significant difference in the expression of verbascoside or aucubin in the leaf tissue across plantain grown in the different soils with the only notable deviation from trend being these metabolites in roots from plants grown in the Waikato soils, where expression was lower in the gley than the allophanic (Supplementary Figure 5).

The phenylpropanoid and iridoid glycosides such as plantamajoside, plantainoside C, asperuloside, and dumuloside were expressed at varying concentrations in the leaf and root material but were low compared with verbascoside and aucubin (data not shown). Of these compounds, dumuloside had a strong influence in defining differences in plantain chemistry in leaf tissue relative to soil type, in addition to plantagoguanidinic acid and a related derivative. This influence was mainly associated with plantain grown in the gley soil and to a lesser extent the allophanic soil (Figure 4). Within the root tissue, 10-benzoylcatalpol was another compound that appears more associated with plantain grown in gley soil (Figure 5).

**Figure 4:**
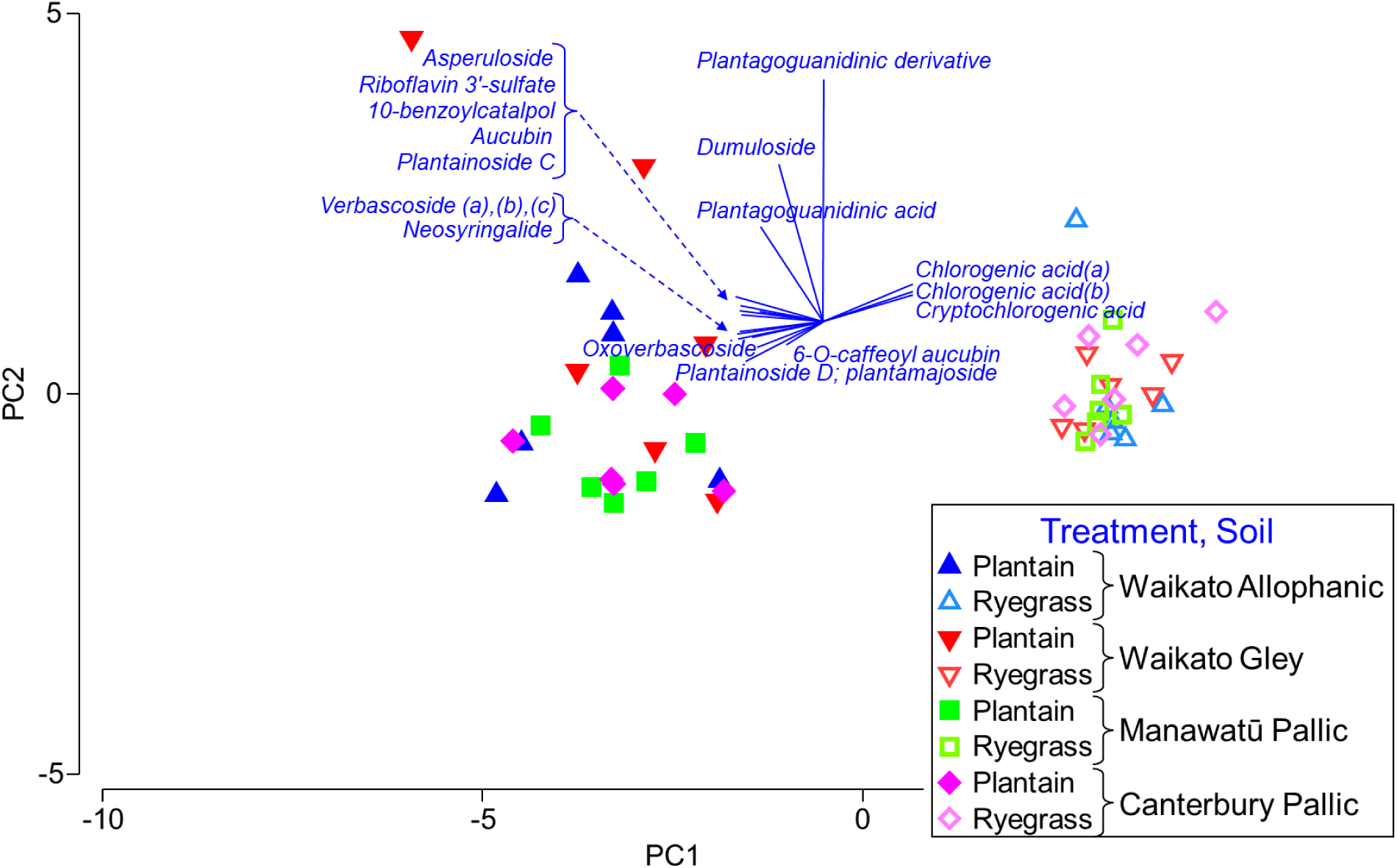
Leaf metabolite expression for plantain cultivar ‘Agritonic’ and ryegrass cultivar ‘One50’ when grown in different soil types. 71% of variation is captured by the PC1 axis and 8.2% by PC2. The PC3 axis captured a further 5.5%. The blue vector fan indicates increasing concentration of each metabolite from the centre out, with length and the direction being an indicator of ‘influence’ that each metabolite has with respect to where data points are positioned in the plot relative to each other.

**Figure 5:**
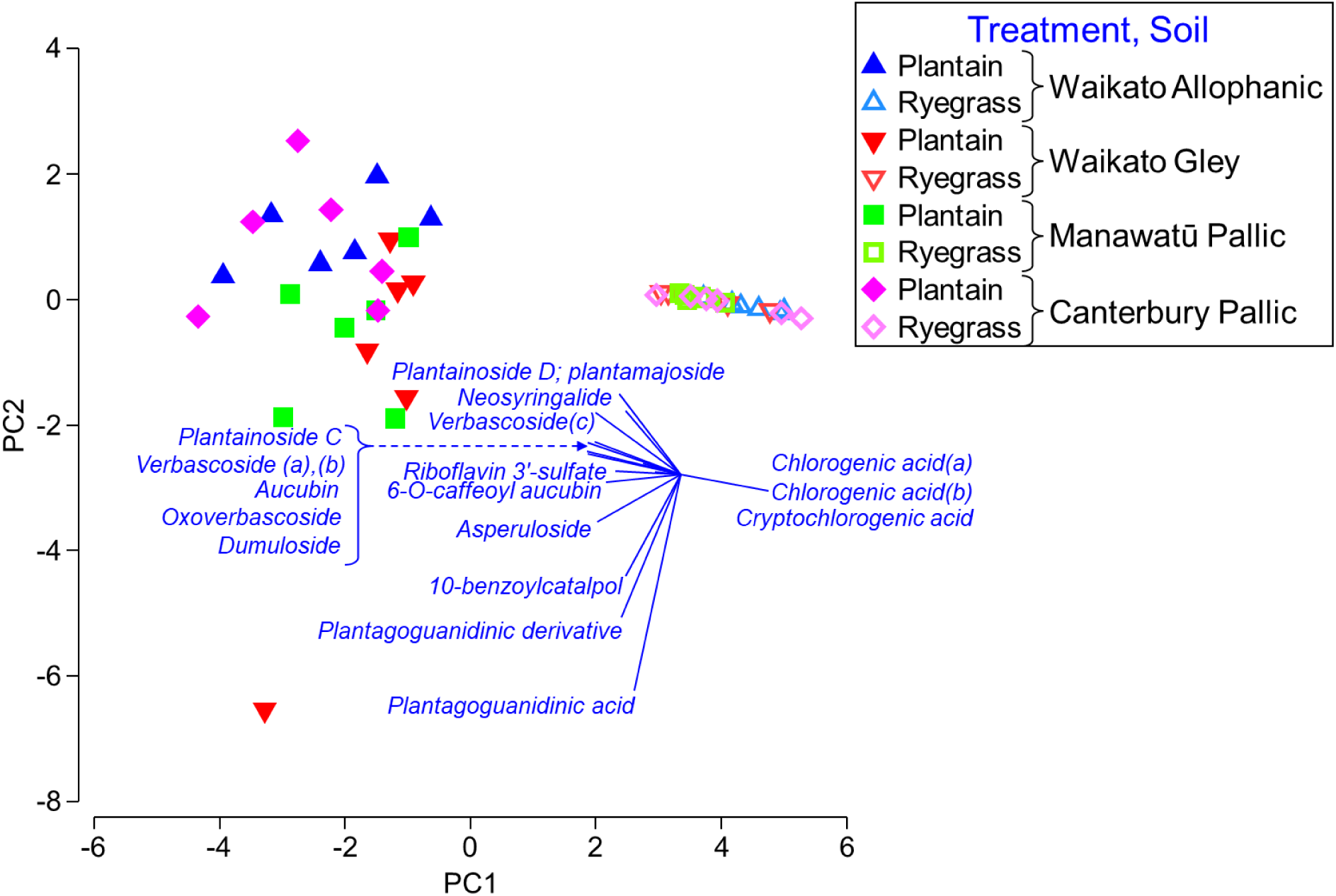
Root metabolite expression for plantain cultivar ‘Agritonic’ and ryegrass cultivar ‘One50’ when grown in different soil types. 67.9% of variation is captured by the PC1 axis and 10.7% by PC2. The PC3 axis captured a further 6.7%. The blue vector fan indicates increasing concentration of each metabolite from the centre out, with length and the direction being an indicator of ‘influence’ that each metabolite has with respect to where data points are positioned in the plot relative to each other.

All other chemistry had minor influence with chlorogenic acid and related compounds defining ryegrass relative to plantain in both leaf and root tissue, irrespective of soil type, as per hydroponically grown plants. A weak visual trend of increasing concentrations of plantamajoside, neosyringalide, verbascoside, aucubin, dumuloside, riboflavin 3-sulphate and asperuloside was observed in plantain root chemistry relative to soil type following the order Waikato gley<Manuwatū pallic<Waikato allophanic≈ Canterbury pallic, but variation was such that there was no significant statistical support (Figure 5).

A wide array of compounds was detected in the rhizosphere soil, some of which were the same as those detected in leaves and roots such as aucubin. Of the 48 key compounds that looked to be associated with lowered PNR, 19% were defined as natural organic matter (NOM) with associated iron, 33% were unknown features defined only by molecular mass and 13% were unnamed features defined by their broad chemical class, for example, ‘polar sulfate’. These groupings, specifically those that associated together, include variants of the same compounds such as isomers. Variation in metabolite concentration was large, covering up to five orders of magnitude for compounds measured under plantain and up to four orders of magnitude for those under ryegrass - subsequently there was no statistical significance for any compounds measured. For example, the highest concentration of aucubin measured in soil under plantain was 7.45 ug g^-1^ (catechin equivalent), similar in magnitude to concentrations measured within the roots. However, variation covered five orders of magnitude across treatments and even within replicates from the same soil type.

The broad picture was illustrated using a heat map (Figure 6), with metabolites organised via an index of association and sample groupings constrained by soil type. Variables were standardised by total and with square root data transformation. There were several shared compounds that differed in concentration between treatments (plantain, ryegrass, fallow) but there were also treatment specific (plantain) compounds such as aucubigenin. Other detectable compounds in the soil that that were identified as being BNI associated within the root tissue from the hydroponic experiments were asperuloside, plantagoguanidinic acid, a plantagoguanodinic acid derivative, geniposidic acid, 10-benzoylcatalpol, a related aucubin derivative defined as 6-O-caffeoyl aucubin, and sulfate containing organic complexes. It is unknown if the latter compounds have any relationship to the riboflavin sulfates identified in the hydroponic experiments. Some of these compounds did not appear among the 25 compounds that contributed most to the index of association discriminating samples from one another as presented in Figure 6.

**Figure 6:**
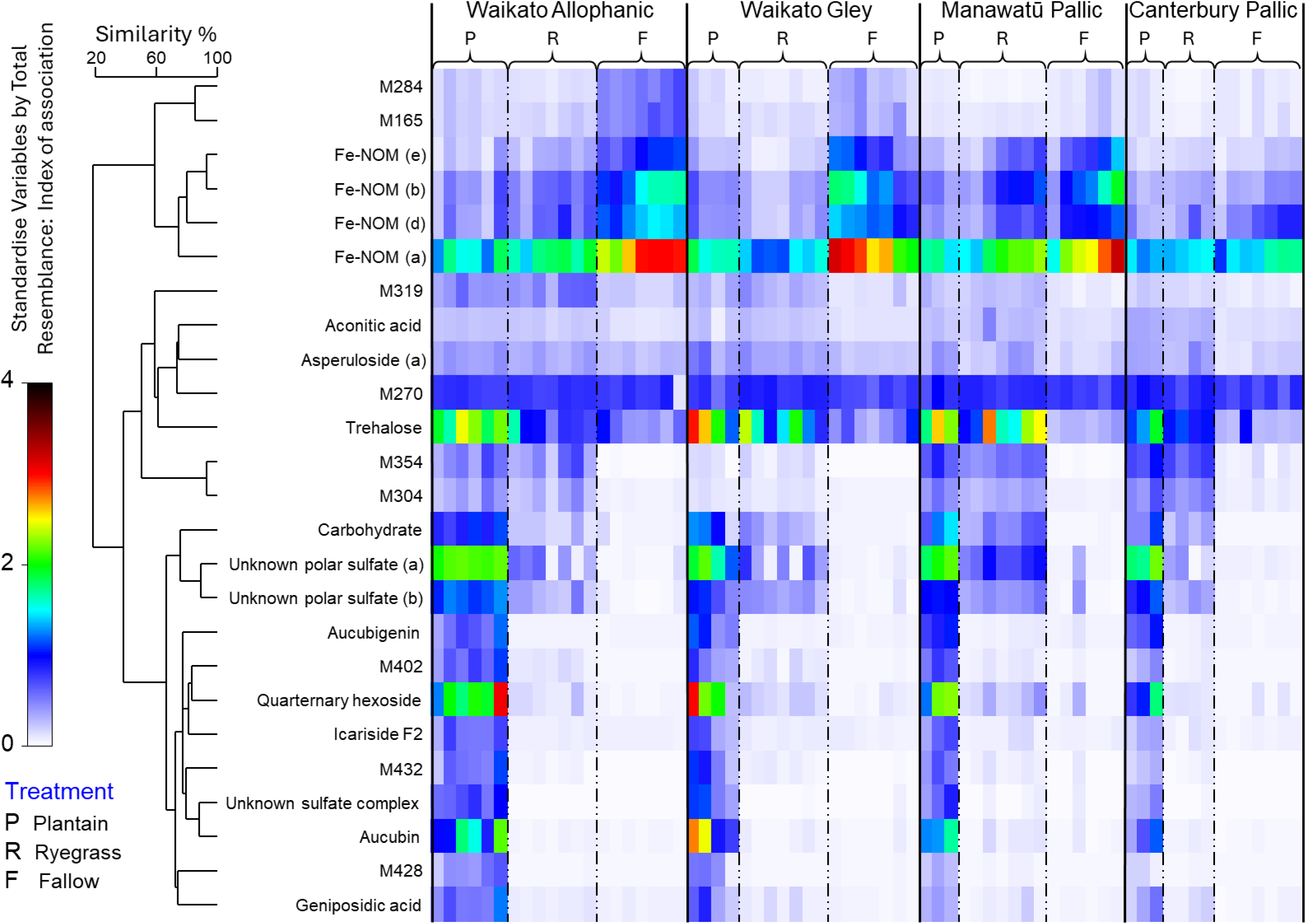
Heat map of metabolites extracted from replicates of rhizosphere soil samples from around the roots of ‘Agritonic’ plantain (P) and ‘One50’ ryegrass (R) along with samples of fallow soil (F). The heat map is organised via an index of association and sample groupings were constrained by soil type. Metabolite data was standardised by total and with a square root data transformation to encompass wide variability. Replicates with indecipherable or no metabolites detected were removed. A total of 48 metabolites were resolved and this heatmap represents the 25 metabolites that best discriminate between samples and treatments. M### designations represent metabolites by their molecular mass if they could not be identified, while Fe-NOM indicates undefined natural organic matter compounds with associated iron. In some cases, only broad chemical features could be ascertained e.g. ‘unknown sulfate complex’.

### Microbial community ecology and assessment of nitrifying bacteria in root associated soil

Higher microbial biomass-N (Table 1) in soil did not appear to be related to higher microbial diversity, rather it was related to greater variability. Chao1 measures of the alpha-diversity within the bacterial community were extremely variable, especially for the Manuwatū Pallic soil, and because of this, significant differences (*p*<0.05) were not observed between treatments or soil types, despite the narrower and lower Chao1 range observed across the replicates in the Waikato allophanic plantain treatment (Supplementary Figure 6). Shannon index measures representing the number of different species and relative abundance were also very similar and lacked statistical significance (Supplementary Figure 6). Analysis of the beta-diversity (between sample differences) by Bray Curtis similarity with square root data transformation, plotted using non-metric Multi-Dimensional Scaling (nMDS), showed a strong delineation of samples by soil type, with the greatest spread among the Waikato Allophanic samples (Figure 7). PERMANOVA supported the difference among soil bacterial communities but within soil type there was no treatment effect, nor interaction, and pairwise tests also indicated no treatment differences. (Table 2).

**Figure 7:**
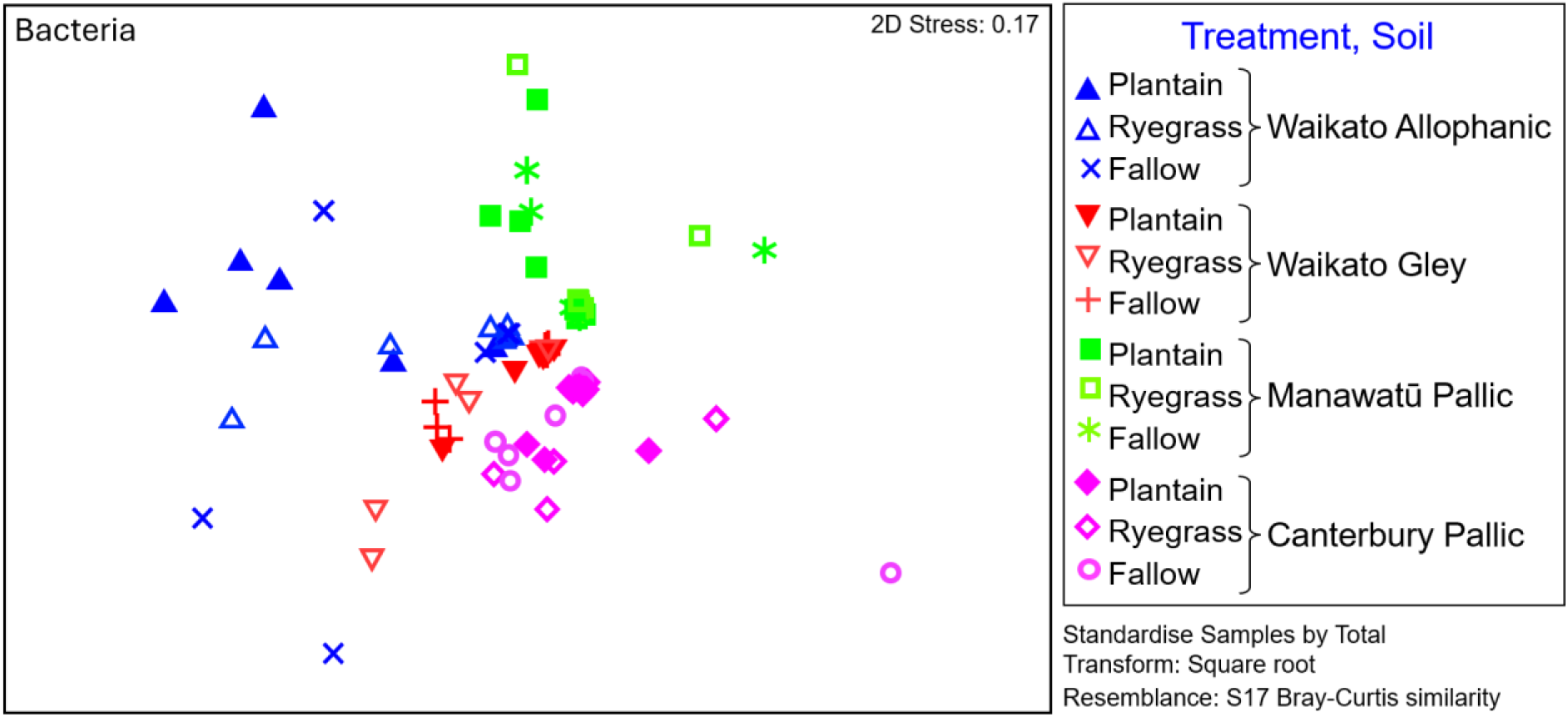
MDS plot of the Bray-Curtis similarities between bacterial microbial communities from four different soils under fallow, plantain or ryegrass treatments. Each soil x treatment combination had seven replicates. The 2D Stress value of 0.17 corresponds to a reasonable 2D representation that should be paired with other statistical testing to support any interpretations made.

**Table 2:** P-values returned from PERMANOVA main tests and pairwise tests investigating the relationships within bacterial or archaeal microbiome data. Each soil x treatment combination had seven replicates. Bold italics indicates significance at *p*<0.05; non-bold italics indicates marginal significance at *p*<0.1.

|  | Permutational p-values |  |
| --- | --- | --- |
|  | *Bacterial communities* | *Archaeal communities* |
| **Main test, all data** |  |  |
| Soil | ***0.001*** | ***0.001*** |
| Treatment | 0.76 | ***0.01*** |
| Soil x Treatment | 0.59 | ***0.011*** |
| **Pairwise tests, treatment** |  |  |
| Plantain, Fallow | 0.695 | ***0.03*** |
| Ryegrass, Fallow | 0.633 | 0.192 |
| Plantain, Ryegrass | 0.643 | ***0.037*** |
| **Pairwise tests, interaction: soil x treatment** |  |  |
| <u>*Waikato Allophanic*</u> |  |  |
| Plantain, Fallow | 0.241 | 0.267 |
| Ryegrass, Fallow | 0.726 | 0.214 |
| Plantain, Ryegrass | 0.345 | 0.169 |
| <u>*Waikato Gley*</u> |  |  |
| Plantain, Fallow | 0.464 | ***0.01*** |
| Ryegrass, Fallow | 0.453 | 0.395 |
| Plantain, Ryegrass | 0.177 | ***0.003*** |
| <u>*Manuwatu Pallic*</u> |  |  |
| Plantain, Fallow | 0.654 | *0.083* |
| Ryegrass, Fallow | 0.502 | *0.079* |
| Plantain, Ryegrass | 0.307 | 0.474 |
| <u>*Canterbury Pallic*</u> |  |  |
| Plantain, Fallow | 0.329 | 0.975 |
| Ryegrass, Fallow | 0.521 | 0.443 |
| Plantain, Ryegrass | 0.538 | 0.17 |

The *Nitrosomonadaceae* family of bacteria, which include the ammonia-oxidising bacteria (AOB), was not among top 20 contributors to the differences in bacterial community structure in any soil or treatment (Supplementary Figure 7). However, bacteria in the *Nitrobacteraceae* family that includes bacteria that catalyse the second nitrite-oxidising step in nitrification (nitrite oxidising bacteria - NOB) do appear and had greater relative abundance in both the Waikato soils compared to the pallic soils.

The relative abundances of the microbial families to which known nitrifying bacteria belong are summarised in Supplementary Table 6. The *Nitrobacteraceae* dominate, with abundance following allophanic>gley>pallic, and the plantain treatment generally had higher abundance of this family compared to ryegrass and fallow. The relative abundance of the *Nitrosomonodaceae* in the Waikato allophanic soil was highest in the fallow treatment and lowest in the plantain. Note that these organisms form a very small part of the microbial community overall and errors were not calculated for this relative abundance data.

The alpha- and beta-diversity measures for the archaeal microbial community indicated that there were significant differences in the numbers of observed species and their abundances (Supplementary Figure 8). The Chao1 indices indicated significant differences (*p*=0.0025) in the number of different species observed while the Shannon indices show significant differences (*p*=0.0056) in the richness and abundance of archaea, particularly between the fallow soils. Plotting via nMDS (Bray Curtis similarity with square root data transformation) showed a clear separation between archaeal communities based on soil type, with the Canterbury Pallic soil having very little dissimilarity between replicates or treatments (Figure 8).

**Figure 8:**
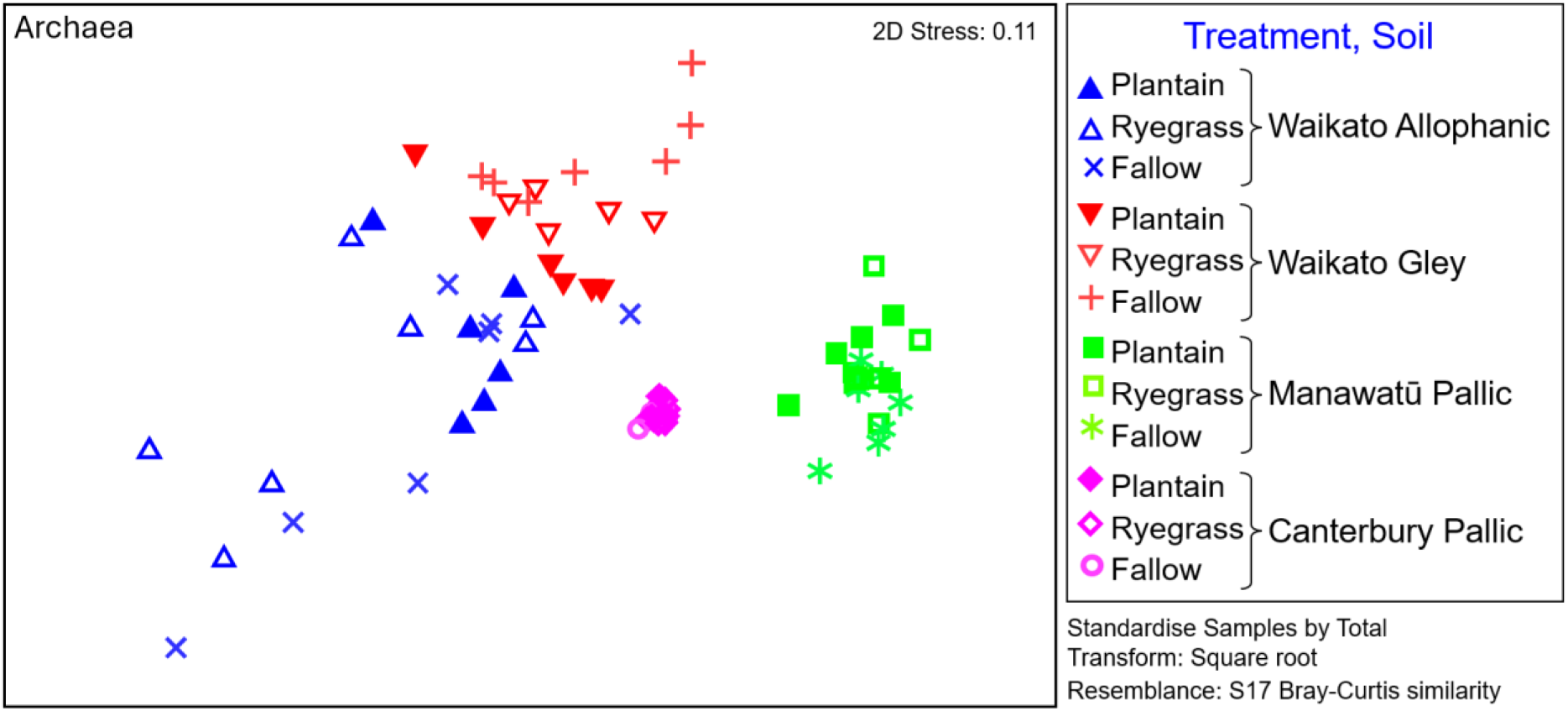
MDS plot of the Bray-Curtis similarities between archaeal microbial communities from four different soil types under fallow, plantain or ryegrass treatments. Each soil x treatment combination had seven replicates. The 2D Stress value of 0.11 corresponds to a good 2D ordination with little prospect of misleading interpretations.

PERMANOVA supported a soil effect, treatment effect and an interaction between the two, with pairwise tests suggesting that archaeal communities under plantain were different to those under ryegrass and fallow, but communities under ryegrass were not significantly different to fallow (Table 2). Pairwise tests investigating the interaction effect indicated that the Waikato Gley soil was driving these relationships, with the only other notable differences being between plantain versus fallow and ryegrass versus fallow in the Manuwatū Pallic soil that were almost significant at *p*=0.05 (Table 2).

Microorganisms from the ammonia-oxidising archaeal phylum *Nitrososphaerota* dominated in the fallow soils (Supplementary Table 7) except in the gley, while the presence of plantain was associated with an increase in the abundance for this phylum in all but the Canterbury pallic soil. At the genus level, *Candidatus nitrosocosmicus* was the dominant AOA in all soil types and accounted for ∼30% of the AOA community in all treatments in the pallic soils (Supplementary Figure 9). An increase in the relative abundance of this AOA was observed in both Waikato soils in the presence of plantain compared to fallow, with a doubling in the allophanic (∼12% to 21%) and a smaller change in the gley (∼22% to 27%). There was no obvious difference between the fallow and ryegrass treatments in these soils. The AOA genus *Nitrososphaera* was found only in the allophanic soils with relative abundances of ∼5%, 2% and 1% for the fallow, plantain and ryegrass treatments, respectively. Note that errors were not calculated for this relative abundance data.

Proportionally, based on averaged qPCR Delta Ct values within the phylogenetic domain, the Waikato allophanic soil had greater bacterial *amo*A and lower archaeal *amo*A than the Canterbury pallic soil, while the values for the Manuwatū pallic and Waikato gley soils lay in between (Table 3). A PCA plot of amoA delta Ct values (Euclidean distance) indicated that soil type had a large influence on the proportions of bacterial and archaeal nitrifiers while plant treatments had little impact. Vectors supported the trend that the allophanic soil generally had lower archaeal *amo*A and pallic soils had lower bacterial amoA (Figure 9).

**Figure 9:**
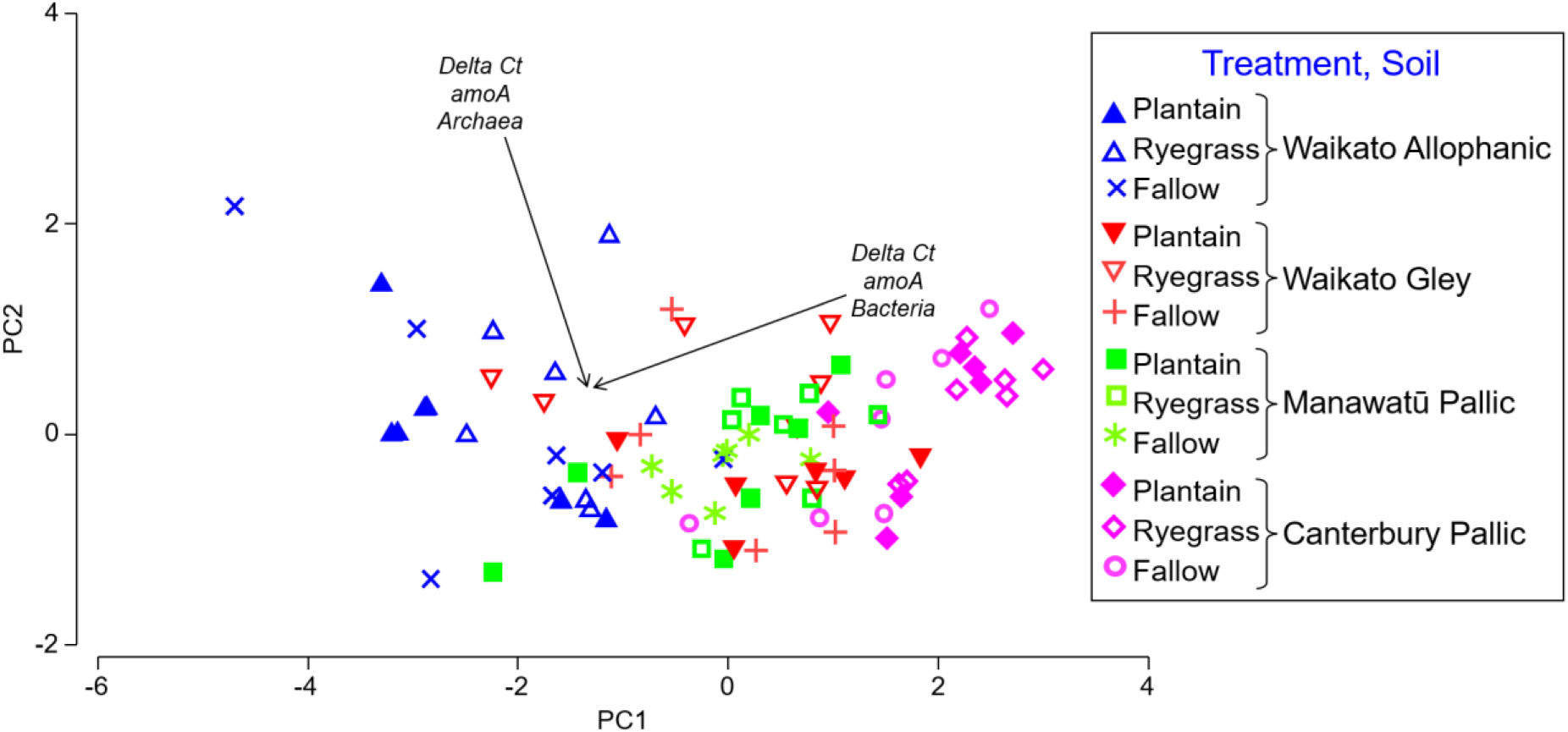
PCA plot for bacterial and archaeal *amo*A Delta Ct replicates from microbial communities across the four soils sourced from three New Zealand regions - Waikato Allophanic, Waikato Gley, Manawatū Pallic and Canterbury Pallic. 84.7% of variation is captured by the PC1 axis and the remaining 15.3% by the PC2 axis. The two vectors indicate increasing levels of the *amo*A gene within the two Kingdoms toward the centre, Bacterial *amo*A (semi-aligned with PC1) has the most influence with respect to where datapoints lie relative to each other on the plot.

**Table 3:** Average Amo Delta Ct values (n = 7) for bacterial and archaeal amoA versus the 16SrRNA gene phyla in four soils sourced from three New Zealand regions – Waikato Allophanic, Waikato Gley, Manawatū Pallic and Canterbury Pallic – that were either left fallow (FL) or planted with plantain (PL) or ryegrass (RG). In general, allophanic soil had lower *amo*A Delta CT for bacteria than pallic soil meaning higher representation of that gene and vice versa for the archaeal *amo*A. Standard deviation for each value is in brackets.

|  | Waikato Allophanic |  |  | Waikato Gley |  |  | Manawātū Pallic |  |  | Canterbury Pallic |  |  |
| --- | --- | --- | --- | --- | --- | --- | --- | --- | --- | --- | --- | --- |
|  | FL | PL | RG | FL | PL | RG | FL | PL | RG | FL | PL | RG |
| Bacterial *amoA* Delta Ct | 7.63<br>(1.21) | 7.21<br>(0.64) | 8.28<br>(0.66) | 9.68<br>(0.83) | 10.00<br>(0.88) | 9.6<br>(1.25) | 9.48<br>(0.50) | 9.33<br>(1.30) | 10.08<br>(0.63) | 10.93<br>(1.09) | 11.57<br>(0.76) | 11.9<br>(0.62) |
| Archaeal *amoA* Delta Ct | 4.85<br>(1.46) | 5.00<br>(0.92) | 4.90<br>(0.90) | 3.86<br>(0.87) | 3.61<br>(0.45) | 4.50<br>(0.81) | 3.82<br>(0.23) | 3.82<br>(0.50) | 3.87<br>(0.49) | 3.69<br>(0.58) | 3.67<br>(0.58) | 3.62<br>(0.39) |

The PERMANOVA main test of *amo*A data supported a soil type effect (*p*=0.001), no clear treatment effect (*p*=0.11) and no strong interactions (*p*=0.47). Pairwise tests for ‘soil type’ were all significant (*p*=0.001), except for the comparison between the Waikato Gley versus Manuwatū Pallic (*p*=0.72) that was also evident in the PCA (Figure 9). Pairwise tests for ‘treatment’ under the main test were almost significant at *p*=0.05 level for fallow versus ryegrass (*p*=0.076), while plantain versus ryegrass was marginal at *p*=0.1 level (*p*=0.097). Among the pairwise tests for the interaction between ‘soil type x treatment’, the only differences that approached significance at the *p*=0.5 level were between plantain versus ryegrass in the Waikato Allophanic soil (*p*=0.057) and between ryegrass versus fallow in the Pallic soils (*p*=0.09 and 0.07 for Manuwatū and Canterbury, respectively).

## Discussion

### Metabolite profiling of hydroponically grown plants and theorised BNI mechanisms

Variation across the six plantain cultivars supports prior research that indicated ‘Agritonic’ performs well with respect to potential BNI (Egan et al. 2025). The bioassay used to assess BNI activity of the six plantain cultivars used in these experiments was variable, but trends allowed selection of cultivars that generally represented ‘high’, ‘medium’ and ‘low’ BNI, but more importantly encompassed a broad range of BNI activity. Previous research has also noted significant variability with this bioassay, which is related to both the physiological state of the nitrifying bacterial species used and the physiological state of the plants themselves when the root exudates were extracted (O’Sullivan et al. 2017b; Kaur-Bhambra et al. 2022; Kolovou et al. 2023). These studies also noted that different nitrifying organisms exhibited different sensitivity and response to the same root exudate preparation.

The initial preliminary screen of the six cultivars grown in hydroponics focussed on identifying metabolite targets that could be used to separate cultivars based on BNI performance, and more specifically distinguish the cultivar ‘Agritonic’ given its generally good BNI activity in bioassays and positive agronomic benefits (Egan et al. 2025 and cultivar information: https://www.agricom.co.nz/products/herbs/plantain/agritonic).

BNI associated with plants grown in the hydroponics system appeared to be due to higher expression of phenylethanoid glycosides. Although the BNI-implicated compound verbascoside (Gardiner et al. 2020) was noted to have the highest concentration in both plantain leaf and root tissue, there was no correlation between verbascoside concentration and the BNI activity ranges defined in this study. Variation in verbascoside with plant age, nutrient supply, temperature and light quality is well documented (Bowers and Stamp 1993; Tamura 2001; Box et al. 2019). This is not to say that verbascoside isn’t involved in BNI, rather the BNI mechanisms are likely more complex and probably involve a wider range of metabolites. For example, secondary hydroxytyrosol compounds in animal urine derived from verbascoside have been shown to be involved with BNI in the soil along with related phenylethanoid glycosides including (R)-ß-hydroxyverbascoside, oxoverbascoside and campneoside (Peterson et al. 2026) (Table 4).

**Table 4:**
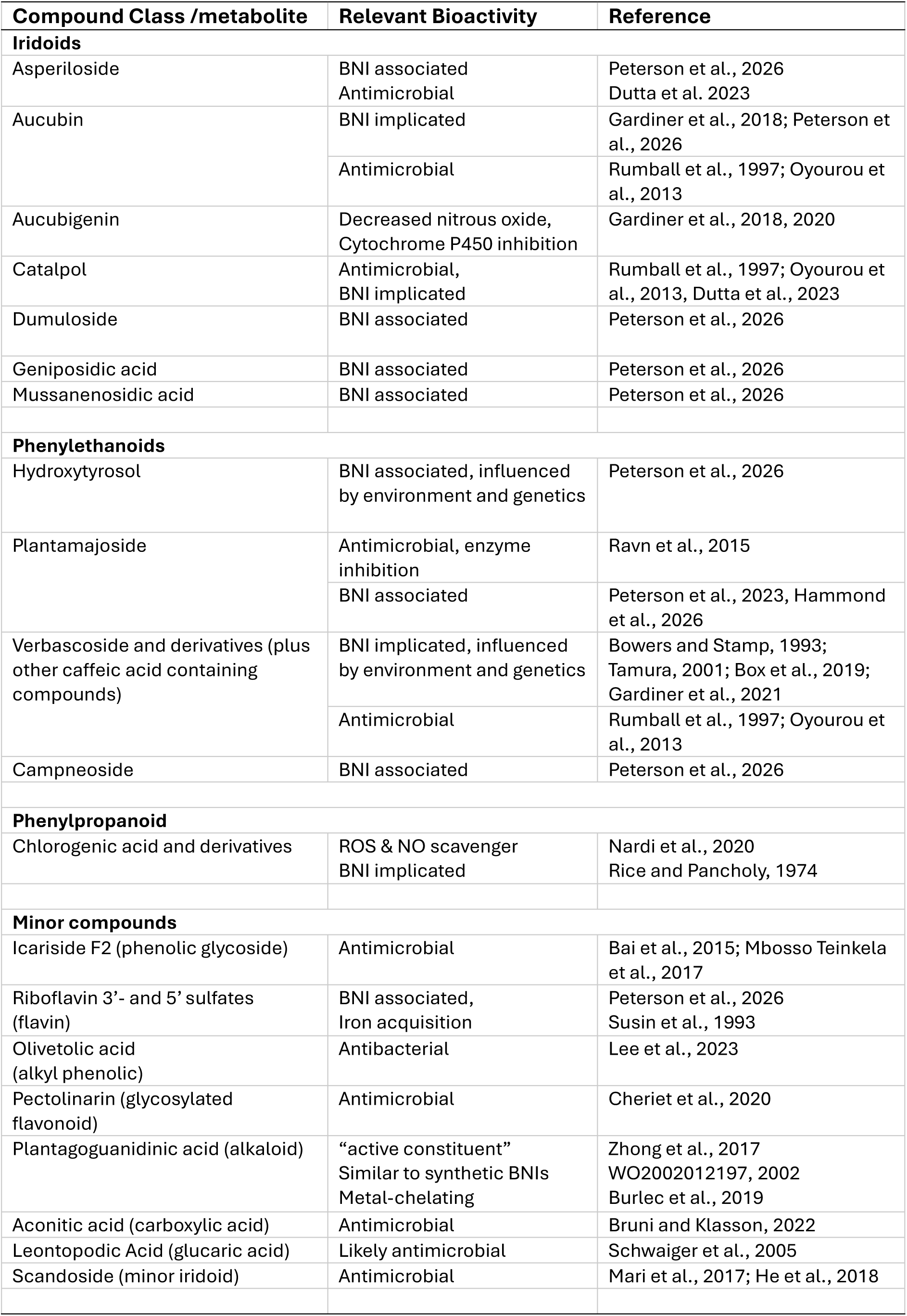
BNI associated plant metabolites detected in this study with reported bioactivities.

Among the plantain leaf tissue phenylethanoid glycosides, plantamajoside was the best correlative BNI biomarker. Plantamajoside has been previously described in several *Plantago* species and is associated with various biological properties such as being an antioxidant, antibiotic, antifungal, cytotoxic, anti-inflammatory, and an enzyme-inhibitory compound (Ravn et al. 2015). In the roots, plantamajoside was not discriminatory with respect to BNI activity ranges but was still a dominant metabolite at concentrations up to 42 mg g^-1^ (phloretic acid equivalent). Even though root concentrations of plantamajoside were not correlated with elevated BNI, it has been implicated with reductions in nitrogen losses from soil (Peterson et al. 2023; Hammond et al. 2026). Peterson et al. (2026) have suggested that BNI is tentatively linked to iridoid glycosides including asperuloside, geniposidic acid, dumuloside, mussanenosidic acid and aucubin (Table 4). All these iridoids were identified in leaf tissue, with the current study identifying additional biomarkers that correlated with BNI activity ranges, including the flavonoid pectolinarin, riboflavin 3’-sulfate and riboflavin -5’-sulfate, and the guanidine alkaloid plantagoguanidinic acid. Of these compounds, aucubin (plus related compounds), asperuloside, plantagoguanidinic acid (plus a derivative) and the two riboflavin sulfates were also detected in the roots and were discriminatory with respect to the BNI activity ranges defined.

Aucubin was present in plantain at concentrations up to 55 mg g^-1^ (phloretic acid equivalent) and is the only metabolite identified in this study that has been further researched with respect to BNI in the field, along with the derivative aucubigenin (Gardiner et al. 2018; Gardiner et al. 2020). In these studies, no robust evidence was presented for nitrification inhibition in the soil, only a lowering of nitrous oxide emissions. Laboratory soil incubations by Dietz et al. (2013) contradict this, as aucubin application to soil resulted in an inhibitory effect on soil nitrification rates. The proposed mechanisms of action for aucubin in the Gardiner et al. (2018) study was direct binding and interaction causing competitive/non-competitive inhibition of ammonia mono-oxygenase (AMO), or oxidation of substrates into products that inactivate AMO enzymatic pathways, as per the mechanisms suggested by Subbarao et al. (2007). Aucubigenin is noted to inhibit cytochrome P-450 and it has been proposed that its structure could allow inhibition of AMO (Gardiner et al. 2020 and references therein).

Asperuloside has not been associated directly with BNI in soil environments. Similar to other iridoid glycosides such as aucubin and catalpol, asperuloside is known to have antimicrobial activity against soil pathogens such as *Fusarium* (Dutta et al. 2023). These types of metabolites could result in indirect BNI via bactericidal and bacteriostatic action, assuming they are exuded into the rhizosphere soil environment.

Plantagoguanidinic acid was first detected in the seeds of *Plantago asiatica* (Goda et al. 2008), has also been found in the seeds of other *Plantago* species and has been described as an “active constituent” within dried ripe seeds of *Plantago asiatica* used in traditional Chinese medicine (Zhou et al. 2013; Zhong et al. 2017). It is a guanidine alkaloid but has not been reported to have any association with BNI or nitrogen cycling processes in soil. The significance of the guanidine alkaloids in the roots and exudates of ‘Agritonic’ is not clear, but the pyrozole ring in plantaguanodinic acid does share some structural similarity to synthetic pyrazole-based nitrification inhibitors in agricultural use, such as the DMP (3,4-dimethyl-1H-pyrazole) and synthetic derivatives (WO2002012197, 2002).

Synthetic pyrazoles act as metal chelators, with this mode of action being one of the proposed inhibition processes thought to reduce the activity of nitrifying organisms (Corrochano-Monsalve et al. 2021). This inhibition could be via chelation of metal cations (Cu^2+/1+^, Fe^2+/3+^ and Zn^2+^) that are obligate cofactors in the metallo-enzymes that catalyse the first two steps of nitrification (Gilch et al. 2010) and likely other metabolic processes (Helmann 2025). Plantagoguanidinic acid has been shown to have metal-chelating activity with the guanidine alkaloid component presenting free N atoms that exhibit Fe-binding capacity (Burlec et al. 2019), meaning that plantagoguanidinic acid and its derivatives have the potential to remove these essential metal cations or reduce their bioavailability.

The ‘riboflavin-sulfate’ metabolites putatively identified in plantain root and leaf biomass also point to chelation as a possible BNI mechanism in this study. Riboflavin-sulfate metabolites allow *Plantago* species to sequester iron (Fe) when presented with it in the Fe(III) form (as in the hydroponic solution in the form of EDTA Fe(III)Na). The importance of these metabolites for Fe-acquisition has been previously reported in sugar beet, where iron deficiency triggers the accumulation of both riboflavin 3- and 5-sulfate (Susin et al. 1993). The need for ‘Fe’ assimilation for *Plantago* growth has also been reported, with varying growth rates dependent on dosing, pH and oxidative conditions (Schmidt and Fühner 1998).

Non-grasses (dicotyledons like *Arabidopsis*, pea and tomato) and plantain use the acidification-reduction transporter-based Strategy I response to acquire Fe (Schmidt et al. 1996; Schmidt and Fühner 1998; Schmidt 1999; López-Millán et al. 2000). On this basis, the plantain root would need to excrete/pump H^+^ ions out to acidify the rhizosphere and maintain conditions to reduce Fe(III) to Fe(II), which is supported at the rhizosphere interface by an exchange linked to phenolics, organic acids (e.g. cinnamic acid) and a ferric-chelate reductase enzyme (Schmidt et al. 1996; Yi and Guerinot 1996; Rodríguez-Celma et al. 2011). It could be postulated that riboflavin-sulfate creates a redox bridge to sequester and transport reduced ‘Fe’ to plant cells while the high caffeoyl phenylethanoid tri-glycoside content in the root (e.g. verbascoside and plantamajoside) could be linked to transportation of cinnamic acids through glycosylation allowing efficient transport under polar conditions within the phloem (Rodríguez-Celma et al. 2011; Rajniak et al. 2018). Plantamajoside may be more efficient at supporting this role due to its differing substitution of a glucose rather than a rhamnose as occurs in verbascoside. Organic acids would support acidification at the root tip.

### Metabolite profiling in soil and implications for N-cycling and theorised BNI mechanisms

The short-term growth (90 days) of plantain and ryegrass in four different soils from three New Zealand regions resulted in a consistent and significant reduction in the potential nitrification rate (PNR) compared with fallow soil controls. There were three plants in each pot, with dense root mats formed and high soil root contact. The metabolites associated with BNI identified in both roots and in the rhizosphere of plantain included: aucubin, aucubigenin, asperiloside, plantagoguanidinic acid, geniposidic acid and 10-benzoylcatalpol. Other metabolites identified were trehalose, aconitic acid and icariside F2 while the broader higher-level chemistries identified included quaternary hexoside, sulfate and natural organic matter-iron complexes.

Interestingly, trehalose is an osmoprotectant produced by soil microbes under drought conditions (Iturriaga et al. 2009) so is probably not plant associated at all. Note that the soil for the PNR assays in this work was air-dried. All the other metabolites detected either have some form of antioxidant activity, including metal chelation and interaction with nitric oxide produced during plant stress, or antimicrobial-like activities. For example, aucubin, and derivatives like catalpol and aucubigenin are antioxidants with elevated expression under plant stress such as drought conditions (Wang et al. 2010; Kartini et al. 2023). From these observations it could be speculated that BNI activity in soils might be higher when plants are stressed, or when plants are relieved from stress and residual metabolites are still present in the soil (e.g. after drought conditions). Differential changes in metabolites related to physiological state, including plant stress, will likely contribute to the variability of BNI/PNR observed. Even plant selection and multispecies pressure in a mixed sward can alter metabolomic profiles (Medina-van Berkum et al. 2025).

Chlorogenic acid, leontopodic acid and 3,5-dicaffeoylquinic acid (isochlorogenic acid) were all expressed in higher amounts in the roots of ryegrass, separating this plant from the plantain cultivars investigated. Chlorogenic acid is a long known BNI-active compound (Rice and Pancholy 1974) and is probably associated with the significant reduction in PNR observed under ryegrass in the pallic soils. Ryegrass was included as a “low BNI” plant, but there is increasing evidence for a wide range of BNI capacity in ryegrasses (O’Sullivan et al. 2017b; Sessoms 2022) making our assumption incorrect. These ryegrass-associated compounds are also recognised as antioxidants known to protect e.g. alpine plants from UV and extreme temperatures (Schwaiger et al. 2005). Like plantain, the effect of the interaction between genetics, environment and management on the expression of BNI associated compounds in ryegrass is unknown.

Melich-3 extraction is a proxy for plant induced metal bioavailability in the rhizosphere. With respect to the main enzymatic metal cofactors required for bacterial nitrification enzymes (Cu and Fe), PNR reduction in plantain soil relative to fallow positively correlated with Melich-3 extractable Cu and Fe when calculated as a proportion of Total Cu and Fe in the soil (R^2^ = 0.88 and 0.97 respectively, Supplementary Figure 10). Ryegrass associated PNR reductions also positively correlated with Cu and Fe (R^2^ = 0.60 and 0.83 respectively). Theoretically, soils with higher pH exhibit lower concentrations of available metals, leading to plants exuding metabolites like organic acids and potential chelators that favourably alter rhizosphere chemistry. Prevalent exudation could conceivably result in elevated trace metal uptake or attenuation by plant metabolites, thereby lowering metal availability for bacterial uptake and use as cofactors in nitrification enzymes, leading to an indirect nitrification inhibition effect. In the context of these experiments, this ‘chelation’ theory could be plausible when comparing e.g. the Waikato gley (pH 5.93) versus the Canterbury pallic soil (pH 6.67) (Supplementary Table 1). Greater exudation of e.g. aucubigenin in a higher pH soil would also support competitive/non-competitive inhibition of AMO by plant metabolites or inactivated AMO enzymatic pathways as per the proposed mechanisms outlined by Subbarao et al. (2007).

Some compounds detected in the soil are more likely derived from plants as they match those detected in plant biomass and are in higher concentration in rhizosphere soil under plantain or ryegrass compared to fallow (Figures 5 and 6). Others such as Fe-NOM compounds are higher in fallow soil compared to rhizosphere soil and are therefore more likely to be soil derived. Assuming Fe-NOM compounds are derived from the soil rather than the plants, this could further support chelation being an indirect BNI mechanism. Root exuded organic acids such as aconitic acid (a citric acid derivative) could lower rhizosphere pH and change redox conditions such that Fe and other metals might be stripped off the natural organic matter (NOM) and transported into the plant, resulting in lower rhizosphere soil concentrations and higher plant concentrations (Janke et al. 2018; Doyama et al. 2021). Unfortunately we did not measure metals in the plant biomass as part of this study, but *Plantago* species are known to bioaccumulate and translocate metals such as iron (Galak and Shehata, 2015).

Of the other named plant metabolites detected in the rhizosphere soil, all have some level of inhibition-related bioactivity (Table 4). Aside from potentially lowering rhizosphere pH, aconitic acid is known for its role as an antifeedant, antimicrobial, and nematicide in plant root exudates (Bruni and Klasson 2022). Icariside F2 is an aromatic glycoside known for anticancer/anti-inflammatory properties (Bai et al. 2015) and has been found in woody plant metabolite extracts that have antimicrobial activity (Mbosso Teinkela et al. 2017). Scandoside is a bioactive iridoid glycoside like catalpol and is found in plants such as *Hedyotis diffusa* (*Scleromitrion diffusum*) and *Paederia scandens*. Scandoside is known for its anti-inflammatory properties, specifically suppression of NF-κB and MAPK signalling pathways, but it is also noted to have potential antimicrobial, antibiofilm, and antitumor activities (Mari et al. ; He et al. 2018). Finally, olivetolic acid is a phenolic acid known via research on cannabinoids where derivatives of this metabolite are known to have antibacterial activity (Lee et al. 2023).

### Rhizosphere soil biology, variation in PNR response, and linkages to phytochemistry

The conditions under which the PNR assays are run, targets a wide spectrum of soil microbes responsible for the conversion of ammonium to nitrate. Although PNR could be considered a blunt instrument, the results from this assay are likely to be more reflective of field conditions, compared to screening BNI in hydroponically grown plantain cultivars using a single organism like *Nitrosospira multiformis* in the *in vitro* bioassay. The PNR assays also captured a snapshot of the overall biogeochemical picture that includes a myriad of metabolic unknowns with respect to the organisms represented by the ∼60% of amplicon sequence variants (ASVs) that returned ‘unassigned’ or ‘undetermined’ at family or lower taxonomic levels in this study.

Similar to Nardi et al (2020), soil type and inherent physicochemical conditions exerted greater influence in these experiments with respect to overall microbial biomass, the proportions of AOA versus AOB, microbial diversity at lower levels of taxonomic classification, and the associated metabolites expressed. Widespread detection of trehalose, a microbially-linked osmoprotectant as mentioned above, also points to the soil having an overarching influence on microbial phenotype expression, with only the archaeal communities seemingly responding to both soil and plant effects (Table 3 - Waikato gley and the Manuwatū pallic soils).

This doesn’t rule out the possibility that plant metabolites have affected alpha diversity within each soil type, just that it is not significant when considering the 5-fold variation observed for metabolites detected in soil from this study (Figure 6). As an example, geniposidic acid - which was present at higher concentrations under plantain - is known to effect significant rhizosphere microbial change (Wang et al. 2025; Zhang et al. 2025), yet irrdoid glycosides like geniposidic acid are often produced in response to pathogen infection, plant-microbe interactions, or drought stress. If the soil was too dry - as implied by the trehalose presence - was the soil physicochemistry the main driver of microbial community structure, the presence of geniposidic acid exuded by the plant in response to stress, or both?

It is possible that there is no metabolite mediated BNI-like response through mechanisms such as those proposed by Subbarao (2007), rather the plants might have simply delivered additional carbon into the soil. This additional carbon could stimulate the microbial community and allow population expansion, leading to immobilisation of N in microbial biomass (Leptin et al. 2021). Immobilisation of N would lead to lower measurable nitrate, giving misleading PNR results. This explanation is plausible for the Waikato allophanic and gley soils under ryegrass which had statistically higher microbial biomass (Table 1) but weakens when considering the PNR for ryegrass soil remained high and was not statistically different to fallow. This suggests nitrogen was not limiting for the PNR assay, nitrate was not sufficiently lowered through microbial biomass immobilisation to affect PNR, and that the PNR response is not misleading. In the pallic soils there was no difference in microbial biomass between plantain, ryegrass and fallow (Table 1), yet PNR was lowered for both plantain and ryegrass supporting the idea that this is a BNI-like effect for both plant species rather than immobilisation of N in expanded microbial biomass lowering perceived PNR.

The potentially mineralisable N (Supplementary Table 2), another chemical measure related to microbial biomass-N and organic-N (Table 1) was at least double for the allophanic soil compared to the gley and pallic soils. These measures suggested the allophanic soil supported a higher microbial biomass, meaning the proportion of the community that can oxidise ammonia was also presumably higher. However, the relative reduction in PNR in the allophanic soil with plantain compared to fallow was approximately one third of that in the pallic soils. The allophanic soil type had high organic carbon and it has been noted that the AOB and AOA community structures alter in response to total organic carbon, favouring AOB (Dai et al. 2018).

Any resulting change in ratio of AOB to AOA would then impact the microbial community sensitivity to potential BNI associated compounds (Kolovou etal. 2023, Kaur-Bhambra et al. 2022) and therefore PNR. Quantitative PCR of the *amo*A gene indicated that the allophanic soil had proportionally higher representation of AOB, so it is conceivable that AOB have a greater contribution to PNR in this soil compared to pallic soils that had proportionally higher AOA. The assumption that bacteria might dominate nitrification processes in this study is also supported by higher representation of the *Nitrobacteraceae* family, members of which catalyse the oxidation of nitrite to nitrate. If the phytochemicals detected from plantain are less effective at inhibiting the activity of AOB relative to AOA, then it might follow that PNR would be lower in the pallic soils compared to the allophanic as observed.

Plantamajoside, verbascoside and derivatives contain a caffeic acid moiety. To add further nuance to the theorised indirect BNI mechanisms postulated above, one proposed mechanism of action for caffeic acid is that it can scavenge reactive oxygen species (ROS) and nitric oxide (NO), a key intermediate in ammonia oxidation to nitrate (Sueishi et al. 2011; Caranto and Lancaster 2017). Removal of NO would then affect the production of nitrate. Chlorogenic acid, the main BNI-associated metabolite implicated in ryegrass, can also scavenge ROS and NO (Nardi et al. 2020). Both caffeic and chlorogenic acid have been observed to have greater inhibition impact on AOA communities than AOB (Kolovou et al. 2023) that could further support the observation of lower PNR in the pallic soils. As alluded to above, these metabolites could also be involved with metal chelation in addition to the riboflavin sulphates identified. This could impact AMO which utilises copper (6 Cu^2+^ and 3 Cu^+^), iron (4 Fe^3+^) and zinc (3 Zn^2+^) as co-factors; and hydroxylamine oxidase a C-type heme protein, which contains 21 heme and 3 non-heme cofactors (Gilch et al. 2010). Future research could investigate the metal adsorption coefficients of these molecules in exudate impacted soil environments with concurrent assessment of the microbial metabolic response with respect to N-cycling and their metallomes.

There are other indirect mechanisms that could impact nitrification rates. Higher PNR in the allophanic soil may be related to the higher amount of carbon allowing heterotrophic microorganisms to outcompete autotrophic nitrifiers as they have higher growth rates and affinity for NH_4_^+^ (Rosswall 1982, Leptin et al. 2021). There will also be indirect metabolic effects of any root exudation that lowers the rhizosphere pH. This could impact nitrification rates as the microbial species involved have optimal pH ranges for both growth and nitrification efficiency (Sahrawat 2008; Amatya et al. 2011; Ayiti and Babalola 2022). It has also been shown that the abundance of both AOA and AOB significantly correlate with the enzyme arylsulfatase, suggesting a relationship between ammonia oxidisers and the availability of organic sulfur compounds (Dai et al. 2018). In this study, organic sulfur in the pallic and gley soils was low (<10 mg S kg ^1^) (Supplementary Table 1) along with lower PNR relative to fallow. With allophanic soil having higher microbial biomass, it could also be that the rate of biodegradation of BNI associated compounds is higher. The high levels of organic matter and allophanic clay could also attenuate the BNI-associated compounds reducing their efficacy as alluded to above. Further work is required to ascertain the validity of these possible impacts on BNI efficacy and closer attention to soil composition aside from N is required to fully ascertain factors that contribute to PNR (and BNI) variability.

### Outcomes and future research

These experiments lie toward the fundamental end of the research spectrum, identifying a collection of candidate BNI-associated/correlated metabolites, before following those through to associated differences in PNR response in four different soil types. Before these metabolites could be carried forward as screening or breeding markers, further investigation is required to establish inhibition activity and to validate the mechanistic theories postulated with respect to the microbes driving the soil nitrogen cycle and the impact of different soil physicochemistries. A key outcome is acknowledging that ribwort plantain cultivar metabolomes can be quite different, meaning no one cultivar will be suitable for all soil and climate conditions with respect to the expectations surrounding plants like plantain in pastoral agroecosystems such as nitrification inhibition and N_2_O emission reductions. Another equally important outcome is that under the conditions of these experiments, perennial ryegrass also exhibited significant ability to lower PNR rates, expressing chlorogenic acid which is a known BNI compound.

Only generic statements can be made regarding the expected performance of plantain cultivars in-field. This is due to variability in soil environments that will have myriad competing chemical, physical and biological factors, along with the variability of the plants themselves and their response to soil and climatic conditions. Given ryegrass elicited similar PNR reductions to plantain in this study, it is also difficult to make definitive statements as to the level of inhibition that might be expected if plantain is included in a mixed species sward and what the full ecological picture might be once animals are also included.

## Conclusions

Our overall hypothesis for this work was that differential expression of BNI associated metabolites delineates BNI capacity and PNR variability across different soil types. This hypothesis was broadly supported but is too simplistic. There are likely many feedbacks between soil physicochemistry, plant physiology and the soil microbiome, which in-turn will influence plant metabolite expression, N-cycling, the soil microbiome and PNR.

The data presented in this work suggests that quantifying only verbascoside, aucubin and catalpol in above ground biomass as the key determinants of BNI activity in plantain is problematic. Other leaf-derived compounds such as plantamajoside and riboflavin-sulphates may be more useful. There are several possible indirect mechanisms that could be attributed to the candidate BNI-associated metabolites hypothesised in this study. One proposed mechanism could be attenuation of trace elements required for the function of key nitrogen cycling enzymes in combination with intermediate N-compounds such as nitric oxide being scavenged by antioxidants. Regarding the roles of the metabolites identified, most literature describes their bioactivity with respect to phytomedicinal properties rather than environmental microbiological and biogeochemical effects. Therefore, any proposed mechanisms postulated in this work require direct experimentation to corroborate theories proposed.

This study also showed that one ryegrass cultivar had potential BNI capability equivalent to plantain depending on soil type. Based on the metabolites identified and the literature, the mechanism(s) of inhibition for ryegrass is likely similar to plantain. In these experiments, the soil chemical environment had the greatest influence over microbiome structure with only the Archaea presenting any evidence of responding to plant exudates found in rhizosphere soil. This likely had a knock-on influence on how the plant derived BNI-associated compounds impacted the overall soil microbiomes, the resulting PNRs observed, and the wide variability.

## Abbreviations

BNI: biological nitrification inhibition
AOA: ammonia oxidising archaea
AOB: ammonia oxidising bacteria
AMO: ammonia monooxygenase
PNR: potential nitrification rate

## Acknowledgements

The authors thank Loreto Hernandez, Sandi Keenan, Emilie Batt, Teri Robson, Richard Gillespie, Chris Dunlop, Vanessa Hampton, Rebekah Tregurtha, Kathryn Lehto, Carlo van den Dijssel, Weiwen Qiu, Mike Cummins, Andrew McLachlan, Heather Jenkins and Duncan Hedderley for their analytical and technical expertise and assistance with the experimental design. The authors also thank Rogerio Cichota, Antonia Miller, Penny Tricker and other collaborators for their review and comments.

## Statements and Declarations

### Author Contributions

***Michelle Peterson:*** Conceptualization, Methodology, Investigation, Formal analysis, Writing – Original Draft. ***Nigel Joyce:*** Conceptualization, Methodology, Investigation, Formal analysis, Writing – Original Draft, Editing and Review. ***John van Klink:*** Methodology, Investigation, Formal analysis, Writing – Review and Editing. ***Preeti Panda:*** Methodology, Investigation, Formal analysis. ***Trish Fraser:*** Conceptualization, Methodology, Project Administration, Supervision. Funding acquisition. ***Craig Anderson:*** Conceptualization, Investigation, Formal analysis, Visualisation, Writing – Original Draft, Writing – Review and Editing, Project administration.

### Data availability

The datasets generated during and/or analysed during the current study are available from the corresponding author on reasonable request. Any requests will require approval by DairyNZ and PGG Wrightson Seeds Ltd.

