## Supplemental tables and figures for "*Plantago lanceolata* and *Lolium perenne* metabolite profiles, their impact on soil microbial community structures and soil biological nitrification inhibition"

**Supplementary Table 1.** Pre-Experiment Sample Site Analysis - Pasture Soil Analysis.

| | pH | EC[A] (μS $cm^{-1}$) | Olsen P[B] (mg P $kg^{-1}$) | Sulfate-S[C] (mg S kg-1) | Organic-S[D] (mg S kg-1) | Allophane (mg kg-1) | P retention[E] (%) | CEC[F] (cmol+ kg-1) | BD[G] (g cm-3) |
|---|---|---|---|---|---|---|---|---|---|
| Waikato Allophanic | 6.15 | 113 | 15 | 49 | 19 | 85,650 | 94.4 | 32.0 | 0.84 |
| Waikato Gley | 5.93 | 101 | 24 | 11 | 9 | 8191 | 53.5 | 21.3 | 1.08 |
| Manawatū Pallic | 6.17 | 87 | 25 | 3 | 7 | 3988 | 29.5 | 17.4 | 1.19 |
| Canterbury Pallic | 6.67 | 113 | 23 | 4 | 5 | 2492 | 19.2 | 13.8 | 1.17 |

[A] Electrical Conductivity; [B] Olsen Phosphorus; [C] Sulfate-Sulfur; [D] Organic-Sulfur; [E] Phosphorus retention; [F] Cation Exchange Capacity; [G] Bulk Density.

**Supplementary Table 2.** Pre-experiment sample site analysis – soil C and N analysis.

| | Total C[A] (%) | Total N[B] (%) | Min N[C] (mg N $kg^{-1}$) | PMN[D] (mg N $kg^{-1}$) |
|---|---|---|---|---|
| Waikato Allophanic | 8.2 | 0.84 | 49 | 221 |
| Waikato Gley | 4.3 | 0.40 | 27 | 123 |
| Manawatū Pallic | 2.8 | 0.27 | 33 | 99 |
| Canterbury Pallic | 2.8 | 0.25 | 33 | 120 |

[A] Total Carbon; [B] Total Nitrogen; [C] Mineral Nitrogen (Ammonium + Nitrite/Nitrate); [D] Potentially Mineralisable Nitrogen.

**Supplementary Table 3.** Pre-experiment sample site analysis – Mehlich 3 extractable soil nutrients.

| | P[A] (mg $L^{-1}$) | K[B] (mg $L^{-1}$) | Ca[C] (mg $L^{-1}$) | Mg[D] (mg $L^{-1}$) | Na[E] (mg $L^{-1}$) | S[F] (mg $L^{-1}$) | Fe[G] (mg $L^{-1}$) | Mn[H] (mg $L^{-1}$) | Zn[I] (mg $L^{-1}$) | Cu[J] (mg $L^{-1}$) | B[K] (mg $L^{-1}$) | Co[L] (mg $L^{-1}$) | Al[M] (mg $L^{-1}$) |
|---|---|---|---|---|---|---|---|---|---|---|---|---|---|
| Waikato Allophanic | 12 | 119 | 1139 | 69 | 19 | 18 | 40 | 30 | 4.3 | 1.4 | 0.32 | <0.1 | 1553 |
| Waikato Gley | 37 | 144 | 1167 | 71 | 15 | 15 | 143 | 135 | 8.4 | 1.1 | 0.38 | 0.3 | 1090 |
| Manawatū Pallic | 49 | 37 | 2160 | 56 | 13 | 10 | 200 | 35 | 0.9 | 1.2 | 0.55 | 0.2 | 873 |
| Canterbury Pallic | 91 | 569 | 1160 | 158 | 22 | 16 | 166 | 25 | 1.6 | 0.6 | 0.55 | 0.3 | 988 |

[A] Phosphorus; [B] Potassium; [C] Calcium; [D] Magnesium; [E] Sodium; [F] Sulfur; [G] Iron; [H] Manganese; [I] Zinc; [J] Copper; [K] Boron; [L] Cobalt; [M] Aluminium.

**Supplementary Table 4.** Pre-experiment sample site analysis – total extractable soil nutrients.

| | S[A] (mg $kg^{-1}$) | Fe[B] (mg $kg^{-1}$) | Zn[C] (mg $kg^{-1}$) | Cu[D] (mg $kg^{-1}$) |
|---|---|---|---|---|
| Waikato Allophanic | 1344 | 19,300 | 127 | 21 |
| Waikato Gley | 583 | 31,000 | 73 | 9 |
| Manawatū Pallic | 444 | 18,500 | 49 | 6 |
| Canterbury Pallic | 276 | 18,100 | 55 | <4 |

[A] Sulfur; [B] Iron; [C] Zinc; [D] Copper

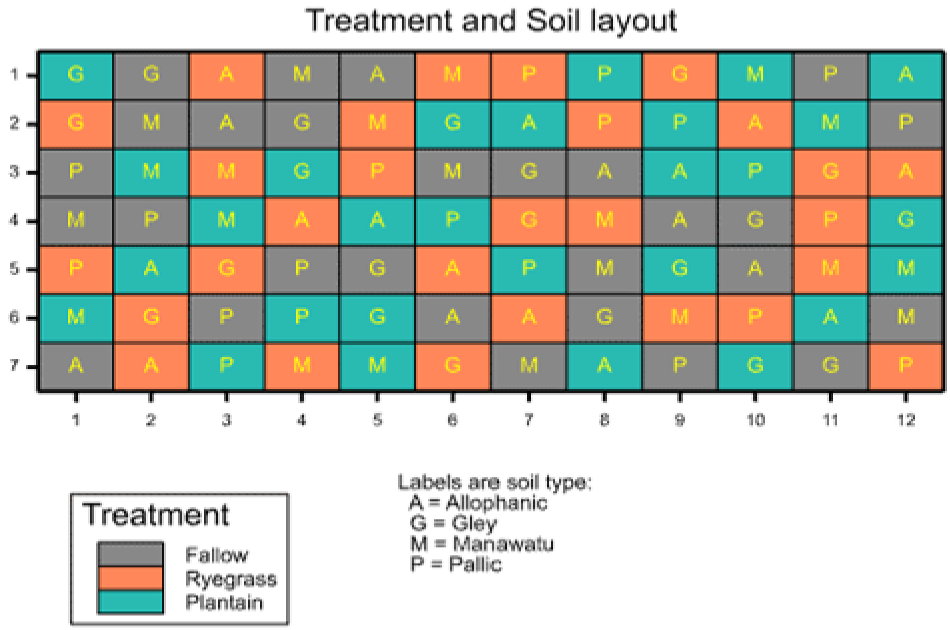


**Supplementary Figure 1.** Spatially adjusted block design for the placement of plantain and ryegrass rhizopots on a sand table.

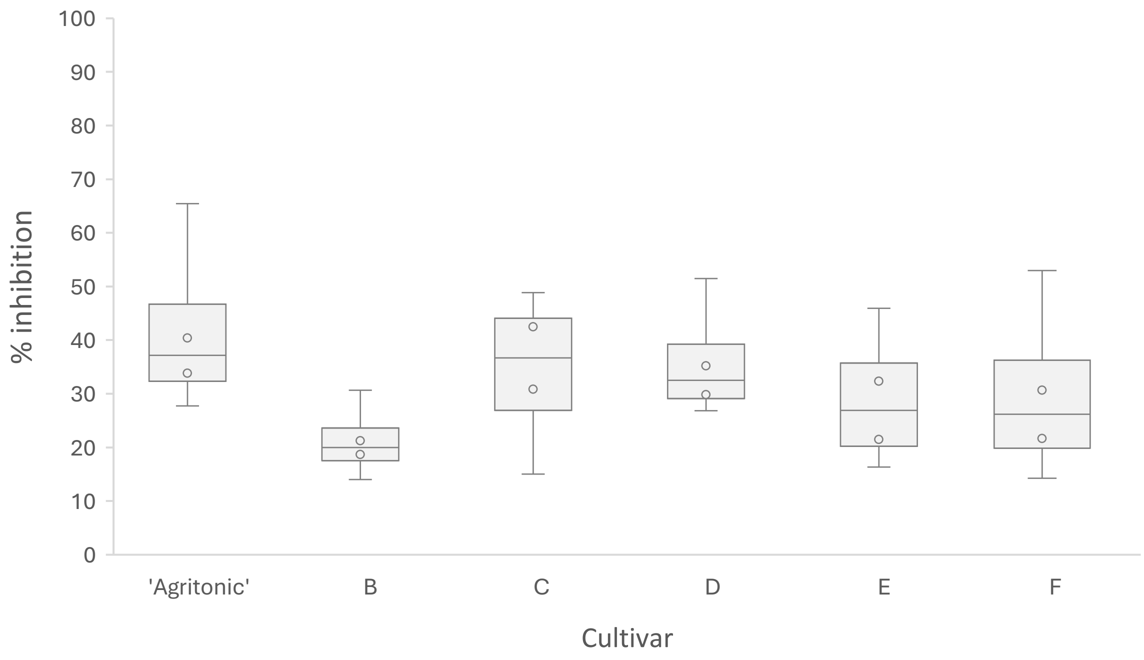


**Supplementary Figure 2.** BNI activity in the root exudates of plantain cultivars, as measured against *Nitrosospira multiformis*, expressed as percentage, calculated from the decrease in nitrification rate in the presence of the root exudate, relative to an uninhibited control. This figure is a permitted reproduction of data published in the Proceedings of IGC 2023, Peterson et al. 2023.

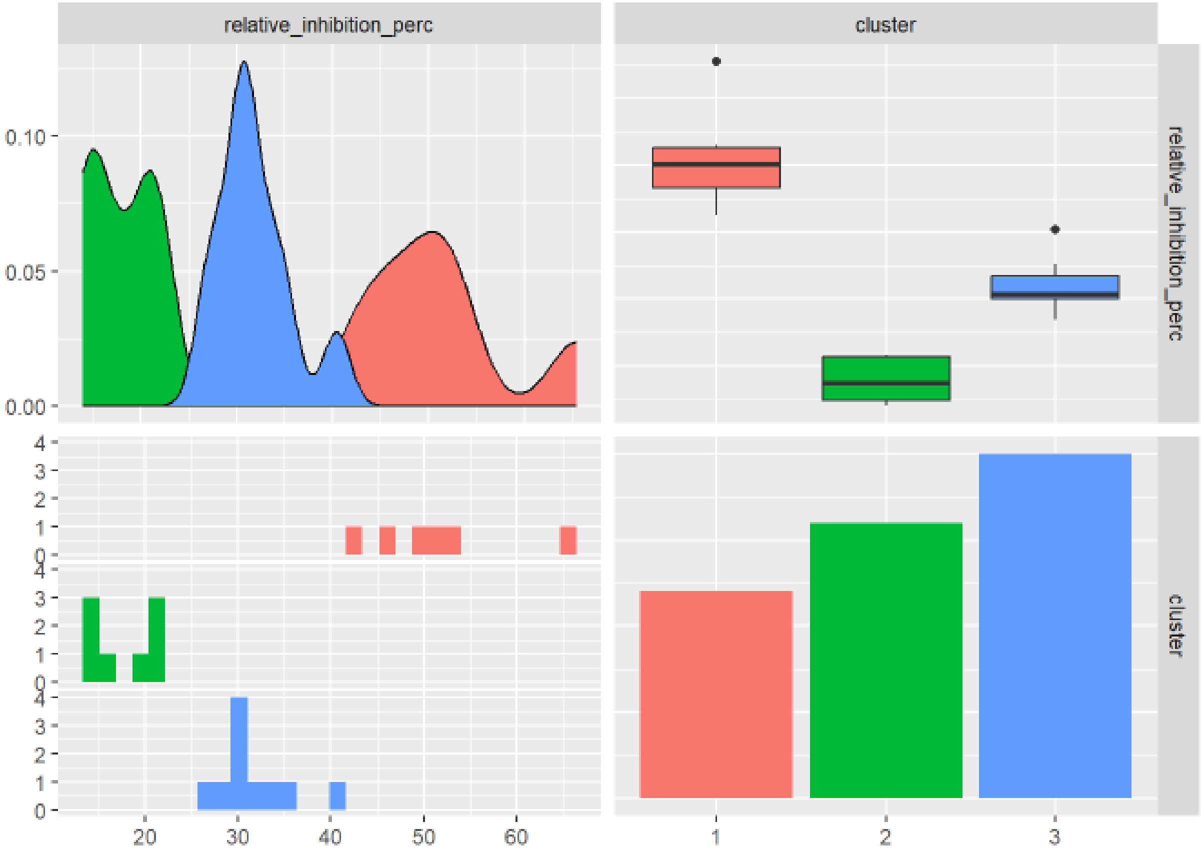


**Supplementary Figure 3.** K-means clustering with 1-8 clusters investigating the total within-cluster sum of squares based on relative BNI percentages reductions compared to uninhibited controls (relative_inhibition_perc). Three clusters were used based on the ‘elbow’ method and were not associated with the cultivars. A Chi-squared test, with null hypothesis “That the cluster numbers are independent of the cultivars” could not be rejected.

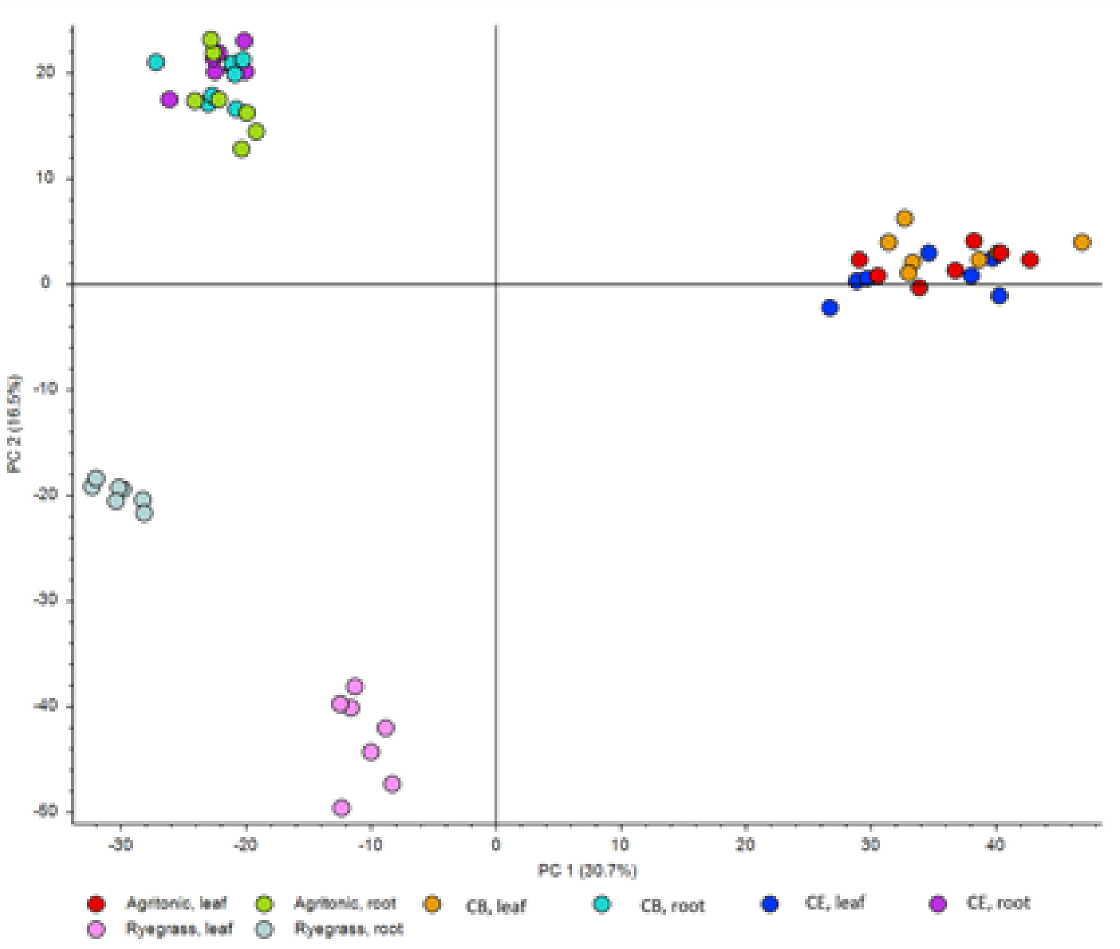


**Supplementary Figure 4.** Principal component (PCA) analysis of liquid chromatography-mass spectrometry (LC–MS) data shows the discrimination of hydroponic plant biomass of leaf and root from three plantain candidates ('Agritonic', Candidate B (CB) and Candidate E (CE)) with reference to ryegrass.

**Supplementary Table 5.** Original ANOVA soil N conditions post plant growth and pre-PNR assay

| | PNR | | PMN | | TON | | MBN * | |
|---|---|---|---|---|---|---|---|---|
| Tests for fixed effects | F | p | F | p | F | p | F | p |
| Soil (3 df) | 700.5 | <.001 | 357.3 | <.001 | 357.3 | <.001 | 168.8 | <.001 |
| Treatment (2 df) | 26.0 | <.001 | 26.8 | <.001 | 26.8 | <.001 | 21.5 | <.001 |
| Soil x Treatment (6 df) | 4.4 | <.001 | 2.4 | 0.024 | 2.4 | 0.024 | 2.6 | 0.029 |

| | MinN | | NOx N | | NH4 N | |
|---|---|---|---|---|---|---|
| Tests for fixed effects | F | p | F | p | F | p |
| Soil (3 df) | 1.7 | 0.184 | 6.2 | <.001 | 3.0 | 0.030 |
| Treatment (2 df) | 155.6 | <.001 | 190.0 | <.001 | 1.6 | 0.198 |
| Soil x Treatment (6 df) | 2.0 | 0.074 | 3.2 | 0.008 | 1.1 | 0.345 |

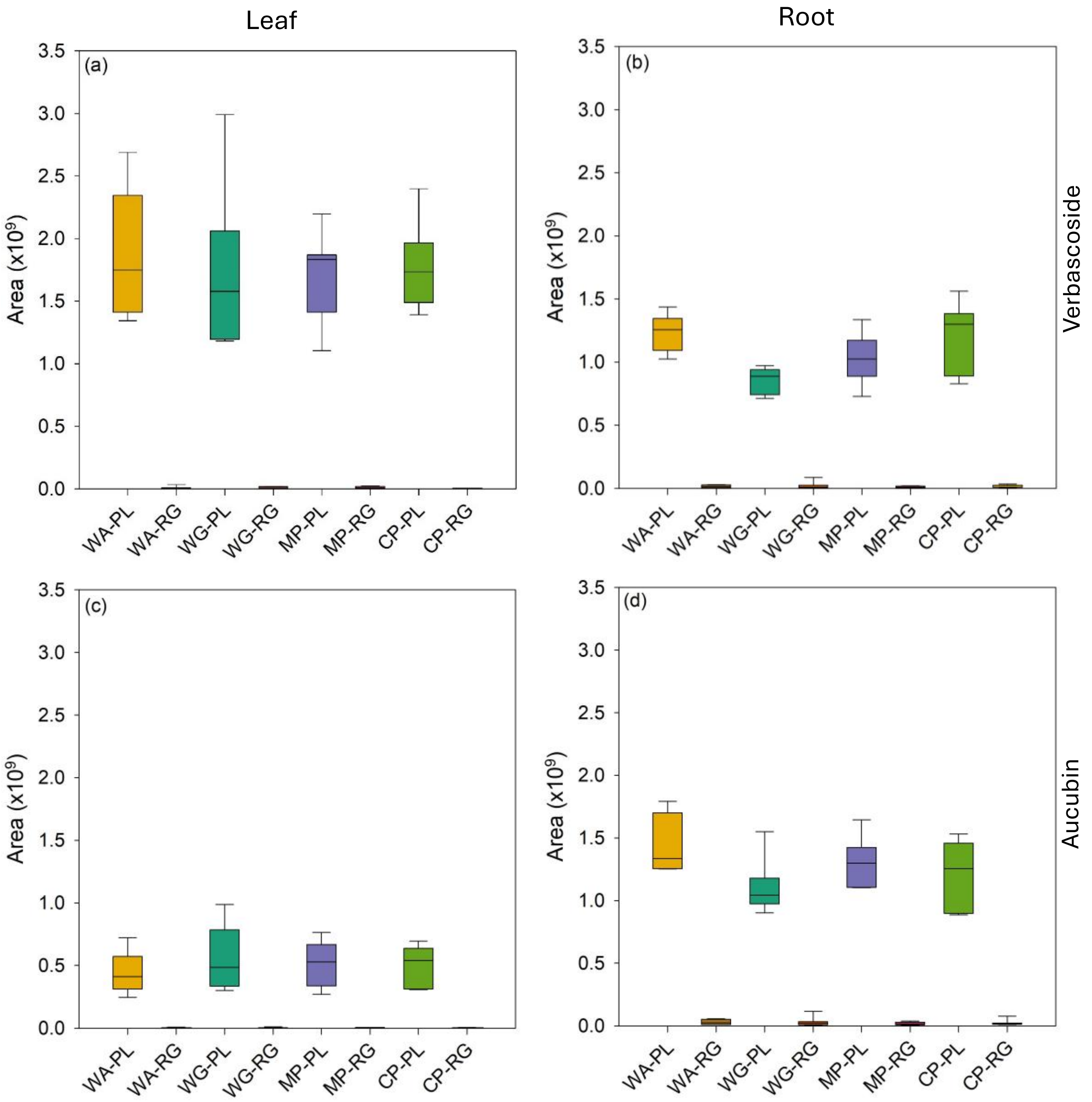


**Supplementary Figure 5.** The abundance (integrated peak areas) of verbascoside in (a) leaf and (b) root; aucubin in (c) leaf and (d) root material of plantain (PL) and ryegrass (RG) grown in four soils sourced from three New Zealand regions – Waikato Allophanic (WA); Waikato Gley (WG), Manawatū Pallic (MP) and Canterbury Pallic (CP).

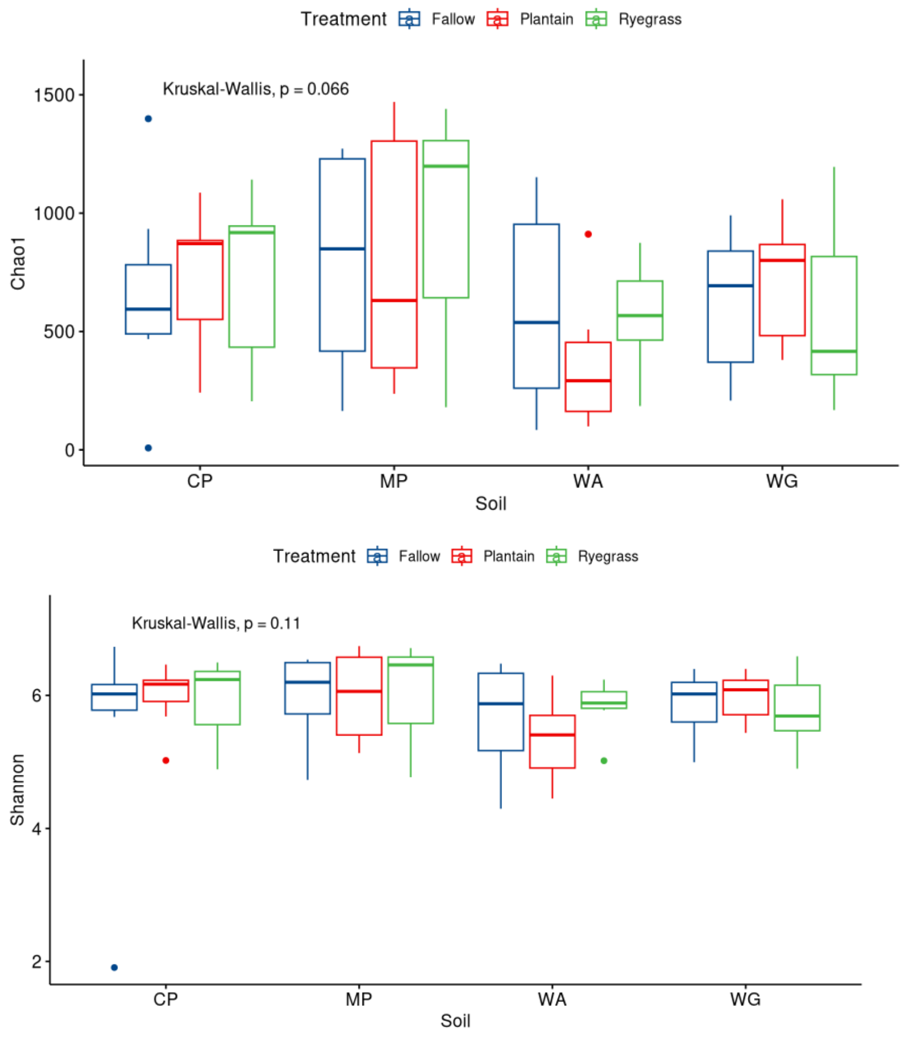


**Supplementary Figure 6.** Bacterial diversity analysis described by Chao1 index, Shannon index and non-metric Multidimensional Scaling (Bray Curtis similarity) in soils sourced from the National Plot Trial sites – Waikato Allophanic (WA); Waikato Gley (WG), Manawatū Pallic (MP) and Canterbury Pallic (CP) – that were either left fallow or planted with plantain or ryegrass.

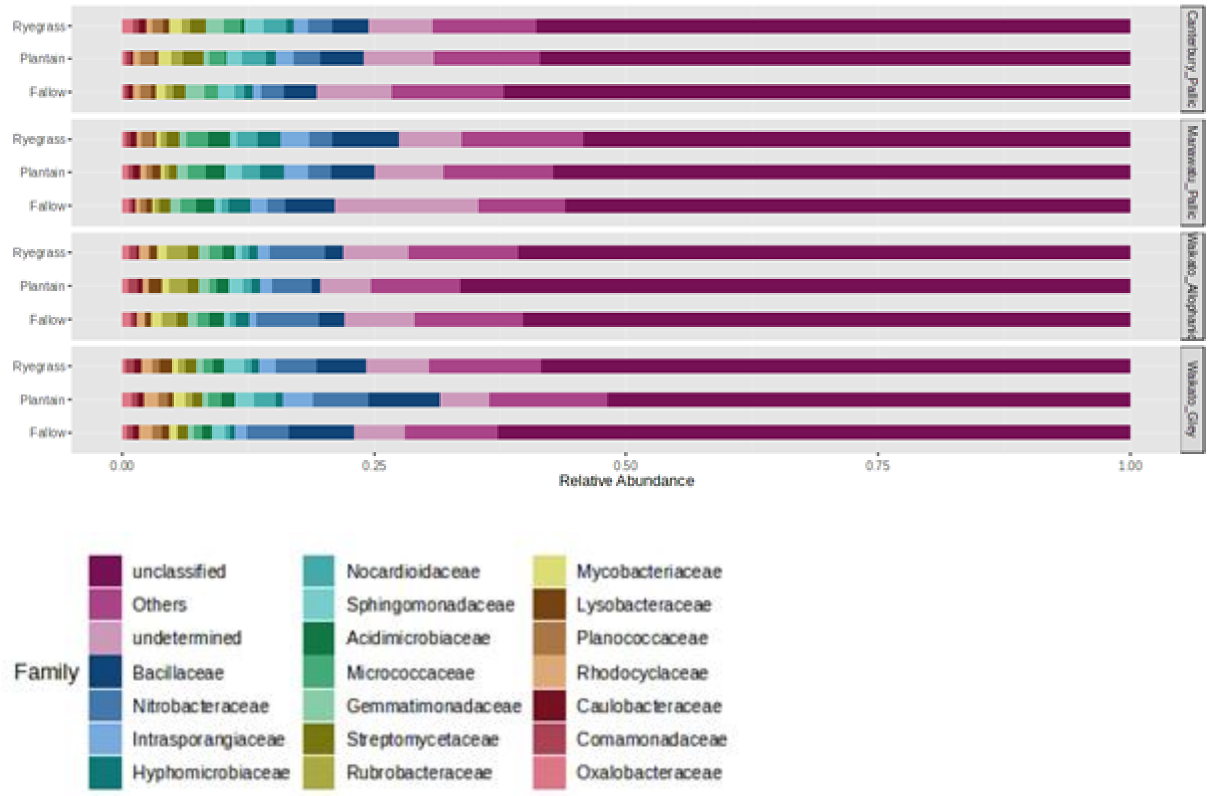


**Supplementary Figure 7.** The top 20 contributors at the bacterial family level that discriminate the soils and their treatments based on relative abundance, normalised to 100%.

**Supplementary Table 6.** The relative abundance (%) of families of known nitrifying bacteria in four soils sourced from three New Zealand regions – Waikato Allophanic, Waikato Gley, Manawatū Pallic and Canterbury Pallic – that were either left fallow (FL) or planted with plantain (PL) or ryegrass (RG). *Nitrobacteraceae* has high representation in the Waikato soils.

| | Waikato Allophanic | | | Waikato Gley | | | Manawatū Pallic | | | Canterbury Pallic | | |
|---|---|---|---|---|---|---|---|---|---|---|---|---|
| | FL | PL | RG | FL | PL | RG | FL | PL | RG | FL | PL | RG |
| *Nitrosomonadaceae* | 0.11 | 0.01 | 0.05 | 0.04 | 0.00 | 0.05 | 0.02 | 0.00 | 0.02 | 0.05 | 0.04 | 0.00 |
| *Nitrobacteraceae* | 6.15 | 3.87 | 5.38 | 4.10 | 5.43 | 3.89 | 1.77 | 2.14 | 2.04 | 2.16 | 2.58 | 2.18 |
| *Nitrospiraceae* | 0.09 | 0.04 | 0.03 | 0.00 | 0.00 | 0.00 | 0.03 | 0.00 | 0.02 | 0.12 | 0.06 | 0.03 |

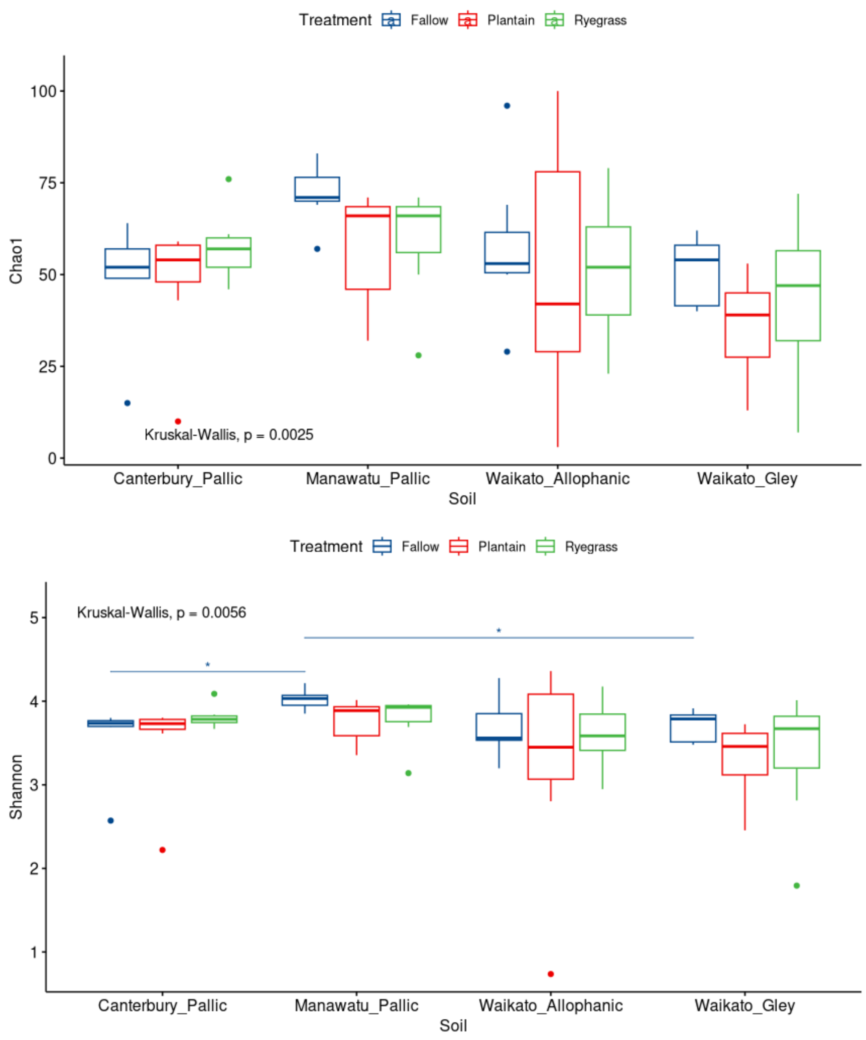


**Supplementary Figure 8.** Archaeal diversity analysis described by Chao1 index, Shannon index and non-metric Multidimensional Scaling (Bray Curtis similarity) in soils sourced from the National Plot Trial sites – Waikato Allophanic (WA); Waikato Gley (WG), Manawatū Pallic (MP) and Canterbury Pallic (CP) – that were either left fallow or planted with plantain or ryegrass.

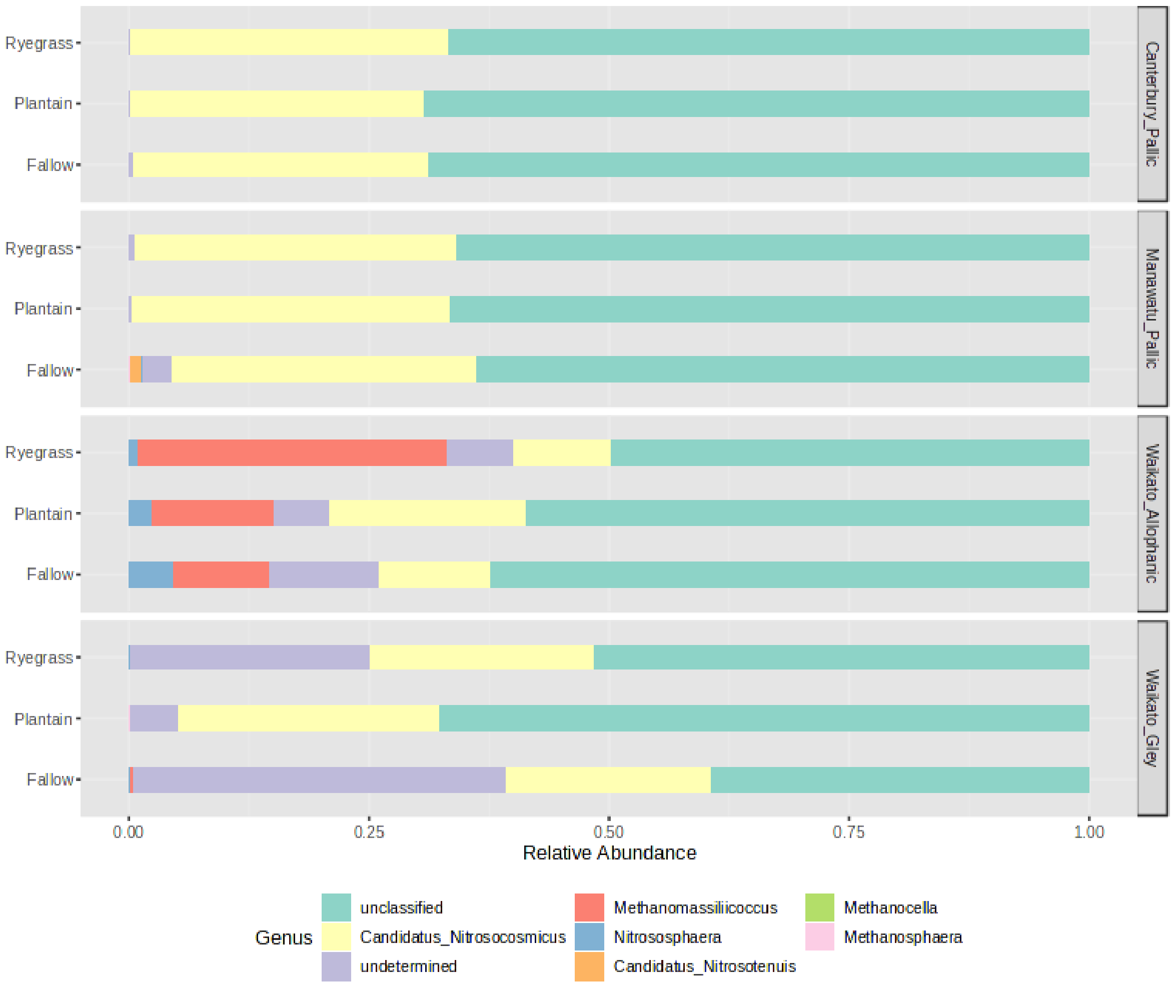


**Supplementary Figure 9.** The archaeal contributors at the genus level that discriminate the soils and their treatments based on relative abundance.

**Supplementary Table 7.** The relative abundance (%) of archaeal phyla in four soils sourced from three New Zealand regions – Waikato Allophanic, Waikato Gley, Manawatū Pallic and Canterbury Pallic – that were either left fallow (FL) or planted with plantain (PL) or ryegrass (RG). *Nitrososphaerota* is the most abundant phyla.

| | **Waikato Allophanic** | | | **Waikato Gley** | | | **Manawatū Pallic** | | | **Canterbury Pallic** | | |
|---|---|---|---|---|---|---|---|---|---|---|---|---|
| | FL | PL | RG | FL | PL | RG | FL | PL | RG | FL | PL | RG |
| *Candidatus Thermoplasmatota* | 11.8 | 12.8 | 34.6 | 1.7 | 0.0 | 0.9 | 0.6 | 0.0 | 0.1 | 0.0 | 0.0 | 0.0 |
| *Euryarchaeota* | 30.1 | 9.2 | 16.5 | 47.2 | 6.8 | 35.0 | 12.6 | 1.5 | 4.2 | 0.8 | 0.0 | 0.9 |
| *Nitrososphaerota* | 52.6 | 70.1 | 43.4 | 44.4 | 82.4 | 53.9 | 79.2 | 89.8 | 86.7 | 89.3 | 89.6 | 88.5 |
| *Thermoproteota* | 5.5 | 7.9 | 5.6 | 6.7 | 10.8 | 10.1 | 7.6 | 8.7 | 9.1 | 9.9 | 10.4 | 10.6 |

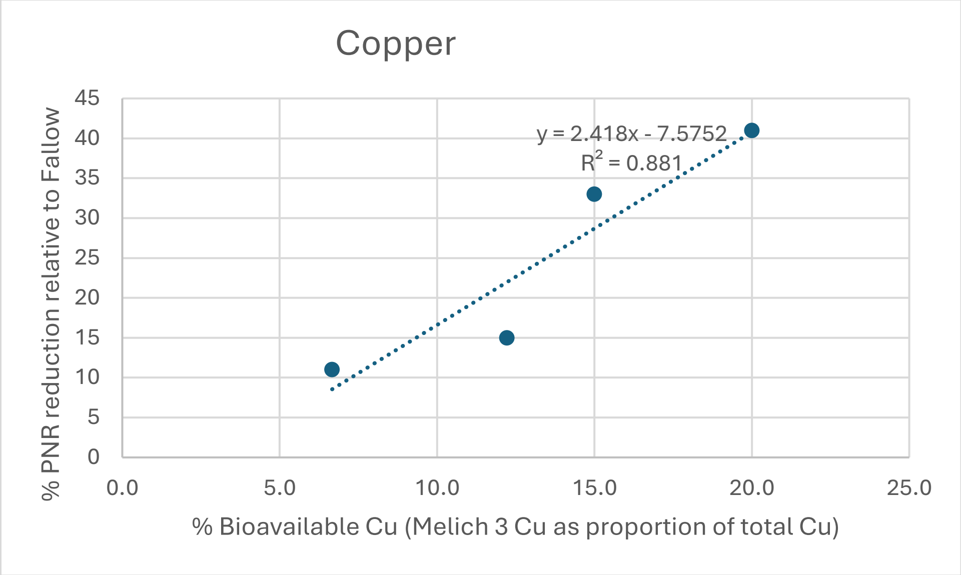


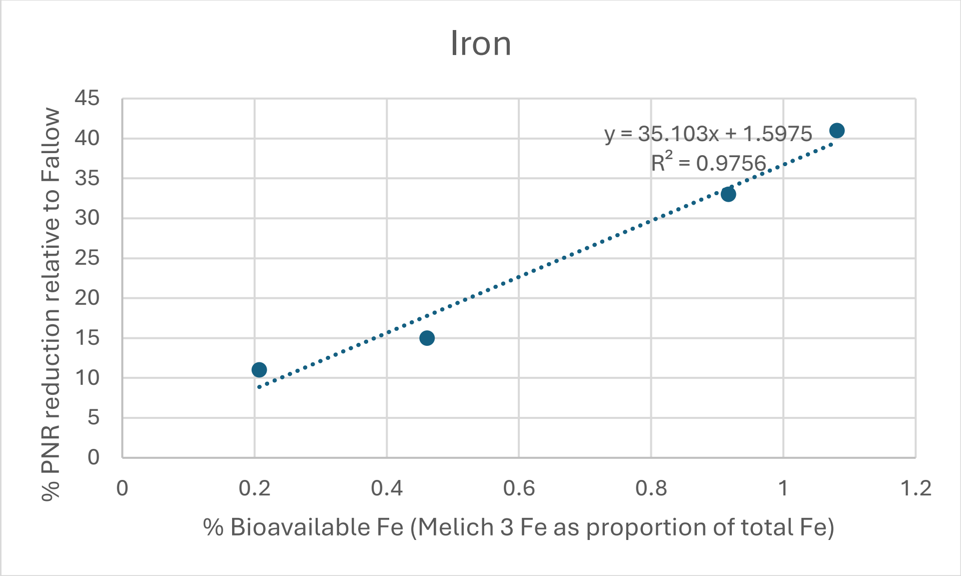


**Supplementary Figure 10.** Regressions between bioavailable metal (Fe or Cu) as a proportion of total versus %PNR reductions relative to fallow soils.
